# ATP-dependent restriction enzyme SauUSI from *Staphylococcus aureus* is also a *bona fide* single strand DNA endonuclease

**DOI:** 10.1101/2022.12.11.519956

**Authors:** Vinayak Sadasivam Tumuluri, Kayarat Saikrishnan

## Abstract

Restriction endonucleases cleave exogenous DNA thus restricting horizontal gene transfer and phage infection of host bacterium. This nucleolytic activity occurs on double-stranded DNA (dsDNA) and is target site specific. Here we report that the Type IV ATP-dependent restriction endonuclease SauUSI from *Staphylococcus aureus* also possesses a hitherto unknown single-stranded DNA (ssDNA) endonuclease activity. We demonstrate that, unlike the dsDNA cleavage activity, ssDNA cleavage by SauUSI does not require divalent cation or ATP hydrolysis and is target-site and DNA methylation-status independent. Furthermore, we show that SauUSI can cut ssDNA gaps, overhangs, bubbles and loops but not ssRNA. The activity is inhibited at higher concentrations of magnesium ion, ATP, and the presence of single strand DNA binding protein. The ssDNA nuclease activity is thus tightly regulated and may protect the host DNA from damage by SauUSI.

## Introduction

Nucleases are enzymes that hydrolyze the phosphodiester bond of nucleic acid backbones and are ubiquitous in all forms of life. They play a pivotal role in various key physiological processes such as replication, transcription and DNA repair apart from regulating transfer of genetic material in the bacterial world (1–6). Foreign genetic material such as plasmids, fragments of double-stranded DNA (dsDNA), double-stranded RNA (dsRNA), single-stranded DNA (ssDNA) or single-stranded RNA (ssRNA) are usually taken up by bacteria by either transformation, transduction or conjugation. More often than not, uptake of foreign DNA can prove detrimental to bacteria, and hence a variety of defense mechanisms to counter the potential catastrophic effects of horizontal gene transfer have evolved. Nucleases that regulate the transfer of genetic material are primarily part of the defense mechanisms of bacteria, such as restriction-modification (RM) systems, clustered regularly interspaced short palindromic repeats associated Cas (CRISPR-Cas) systems and non-specific endo/exonucleases.

RM systems are the most prominent bacterial defense system found in ∼75 % of the sequenced bacterial genomes (7). In comparison, CRISPR-Cas systems are found in ∼40 % of the sequenced bacterial genomes. RM systems are usually composed of two components that act in tandem. The restriction component selectively cleaves foreign DNA and the modification component protects the host DNA from the restriction activity by methylating it at cognate sites. Restriction endonucleases discovered thus far are known to cleave only dsDNA and there has been no known example of a *bona fide* ssDNA endonuclease reported. Though, during the early days of research on restriction endonucleases, there were exciting reports of restriction endonucleases cleaving ssDNA (8–15), they could be explained as arising from cleavage of transient dsDNA formed by the ssDNA (8, 10–12, 16). Here, we demonstrate that the restriction endonuclease SauUSI from *Staphylococcus aureus*, which is a sequence-specific dsDNA shredder (17), is also a *bona fide* non-sequence-specific ssDNA endonuclease.

Previous studies had characterized SauUSI as a modification-dependent Type IV REase that recognizes the target-site 5’-S^5m^CNGS-3’ (S=C/G and N= A/T/G/C) and cleaves dsDNA multiple base pairs away from the target site (17, 18). SauUSI contains a N-terminal PLD nuclease, a SF2 helicase-like ATPase and a C-terminal SRA target recognition domain. Detailed biochemical and structural studies revealed that it functions as a stable dimer in solution with the interface formed by the nuclease domains. Part of its impregnability as a barrier to horizontal gene transfer stems from its unique mechanism of cleavage wherein DNA present between two target sites is shred into multiple fragments (17). This shredding of DNA is brought about by the enzyme’s ability to translocate on dsDNA fueled by the hydrolysis of ATP. In addition to this, SauUSI also cleaves dsDNA with a single target site in an ATP-dependent manner albeit with an efficiency lower than substrates having multiple target-sites.

One of the ways methicillin resistant strains of *S. aureus* are converted to vancomycin resistant ones is attributed to inactivation of the *sauUSI* gene which eventually leads to a manifold increase in horizontal gene transfer efficiency (2, 19). The inactivation of the *sauUSI* gene is hypothesized to occur when the bacterium is challenged by antibiotics where the only means of survival is to take up foreign DNA either from other bacteria (conjugation) or from the environment (natural transformation) (20–23). Presently, vancomycin and combinatorial antibiotics are the last line of defense against methicillin resistant *S. aureus* (MRSA). Hence, the emergence of vancomycin resistant *S. aureus* (VRSA) is a burden on our present medical infrastructure. Understanding the ways in which bacteria evolve antibiotic resistance is of paramount importance to devise novel ways to combat these pathogens.

In this study, we show for the first time that a restriction endonuclease, SauUSI, has the ability to cleave ssDNA, which is DNA sequence-independent and does not require ATP. We also show that ssDNA as short as 25 -base long to very long ones, such as M13mp18 ssDNA, are efficiently cleaved my SauUSI in the absence of nucleotide and without any sequence specificity. We further demonstrate that SauUSI cleaves ssDNA endonucleolytically without a preference for either a 5’ or 3’ free end. We also find that magnesium ion (Mg^2+^), ATP and single strand DNA binding protein (SSB) affect the ssDNA endonucleolytic activity of SauUSI, which may lend protection to the host DNA from the toxicity of the enzyme. This novel view on SauUSI, that is, its ability to cleave both dsDNA and ssDNA, could have major implications in determining the efficiency of HGT and acquisition of antimicrobial resistance.

## Materials and methods

### Purifications of SauUSI

SauUSI and the nuclease-inactive mutant (SauUSI^H119A^) were purified using the protocol described earlier (17, 24). Both the *sauUSI* and *sauUSI* ^*H119A*^ were cloned into a *pHis17* expressed vector containing a C-terminal 6x His-Tag. A three-column purification strategy involving Ni-NTA affinity chromatography, anion-exchange chromatography and size exclusion chromatography, at 4° C was employed to obtain the protein to a high degree of homogeneity.

### DNA Substrates

Sequences of substrates used in this study are mentioned in the supplementary table. The dsDNA substrates used in this study were annealed using commercially synthesized ssDNA oligos. These oligos were annealed by sequentially decreasing the temperature from 95°C to 25°C in steps of 1°C at an interval of 1 minute in each step. The substrates were run on a 12% native PAGE to confirm strand annealing. The ssRNA used in the study was commercially synthesized. The canonical cleavage assays for long dsDNA substrates were generated either by PCR amplification (EcoP15I) or by linearizing plasmids derived from a *dcm+* strain of E. coli (SauUSI, LlaBIII and BstNI).

### Cleavage assays

Initial cleavage assays to test the activity of the enzyme were performed by incubating 250 nM DNA substrate and 200 nM SauUSI with 2 mM ATP in the NEBuffer™ 4 (50 mM potassium acetate, 20 mM Tris-acetate, 10 mM magnesium acetate and 1 mM DTT pH 7.9) at 37° C for 45 minutes. Temperature and incubation time were kept constant for all the assays (unless specified in the results section). Assays that were conducted in the presence of divalent cation chelator were performed in the presence of 16 mM EDTA. Assays that were conducted in the absence of magnesium acetate used a buffer containing 50 mM potassium acetate, 20 mM Tris-acetate pH 7.9 and 1 mM DTT. Subsequent characterization of cleavage assays was performed by keeping the concentration of SauUSI at 50 nM. The reactions were stopped by adding 1X Gel Loading Dye Purple followed by heating at 65 °C for 20 minutes. Cleavage assay with BmrI was performed by incubating DNA substrates with BmrI in the presence of NEBuffer™ 2.1 at 37 °C for 1 hour. The cleavage fragments were resolved on a 12% native PAGE and visualized on a GE Amersham

Typhoon gel scanner after staining with 1x SyBr gold staining solution. Some of the initial assays were visualized on a G-box UV imager. The cleavage assays using different nucleotides (ATP, GTP, CTP, UTP, dATP, ADP and AMP) were performed using the NEBuffer™ 4. The combination cleavage assays were performed with 125 nM 60 bp dsDNA and 250 nM 36-base long ssDNA in NEBuffer™ 4. RNA cleavage assay was performed by incubating 250 nM ssRNA with a concentration gradient of SauUSI in NEBuffer™ 4 at 37° C for 30 minutes. The reaction was stopped by adding 1X formamide loading buffer and heating at 95° C for 10 minutes. The cleavage fragments were resolved on a 15% urea denaturing PAGE and visualized on a GE Amersham Typhoon gel scanner. Densitometric calculations for all the cleavage assays were performed using ImageJ software and plotted using GraphPad Prism.

### ATP-dependent Cleavage assays

The cleavage assay using long dsDNA to test the activity of SauUSI, EcoP15I, LlaBIII and BstNI were performed as described earlier (17, 25, 26). 100 nM of SauUSI was incubated with DNA at 37°C for 45 minutes. 100 nM of LlaBIII or EcoP15I were incubated with DNA and 2 mM ATP at 25°C. BstNI was incubated with DNA at 60°C. All the reactions were stopped using 1X Gel Loading Dye Purple followed by heating at 65 °C for 20 minutes.

### DNA binding study

The DNA used for binding study is a 60 bases ssDNA. 250 nM of ssDNA was incubated with SauUSI^H119A^ on ice for 20 minutes. EMSA was carried out on a 5% native PAGE and the samples were loaded using a 1X STB (sucrose, Tris and bropmophenol blue) loading buffer. The gel was stained using a 1X SyBr gold staining and visualized on a GE Amersham Typhoon gel scanner.

### Cleavage assay using M13mp18 ssDNA

100 nanograms of NEB M13mp18 ssDNA which corresponds to ∼882 nM in a 10 microlitre reaction was incubated with 220 nM, 441 nM and 882 nM of *E. coli* SSB (single-stranded DNA binding protein) to form a SSB:ssDNA molar ratio (R) of 0.25, 0.5 and 1 respectively in a low-salt buffer (2 mM NaCl, 10 mM Tris pH 8 and 0.1 mM EDTA) at 25° C for 2 hours (27, 28). The molarity of M13mp18 ssDNA was calculated assuming a (SSB)_35_ mode of binding and that of *E. coli* SSB was calculated using its tetramer quaternary assembly. The reactions were electrophoresed on a 0.5% agarose gel equilibrated with ethidium bromide in 0.5X TAE buffer and visualized on a E-gel Imager. Following ssDNA-SSB complex formation, 50 nM of SauUSI was added and incubated at 25°C for 30 minutes and the reactions were visualized on a 0.5 % agarose gel equilibrated with ethidium bromide which was electrophoresed as mentioned earlier.

## Results

### SauUSI cleaves ssDNA in an ATP- and target-site independent manner

Previous studies on SauUSI revealed that it has the ability to specifically cleave methylated DNA containing a target-site in an ATP-dependent manner and DNA which is non-methylated is not a substrate for cleavage. Non-methylated dsDNA is refractory to cleavage by SauUSI owing to its inability to stimulate the ATPase activity of the SF2 helicase-like ATPase domain which tightly regulates the nuclease activity. This was exemplified when a 60 bp non-methylated dsDNA was subjected to cleavage by 100 nM SauUSI in the presence of ATP (Figure 1, Lane 2, 3 and 4). As expected, the non-methylated DNA was not cleaved by SauUSI either in the presence or absence of ATP. The 60 bp dsDNA substrate was an annealed product of two 60 base-long ssDNAs.

**Figure 1:**
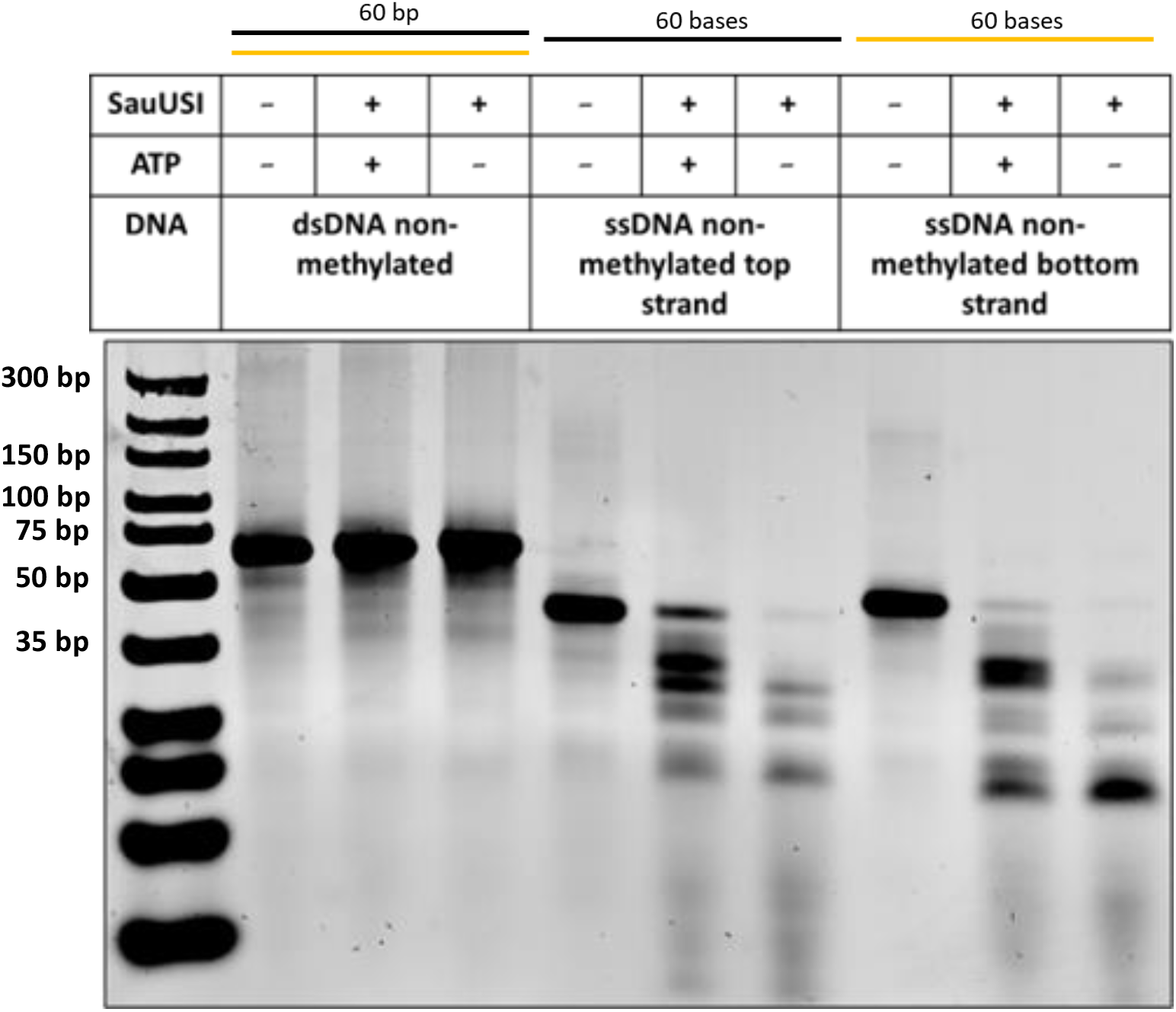
ssDNA cleavage is target site independent and does not require ATP: Schematic of the substrates used for the cleavage assay. Representative 12 % native PAGE gel to study the cleavage assay for dsDNA (non-methylated) and both the ssDNAs (non-methylated top/bottom strands). The dsDNA substrate was generated by annealing the two ssDNAs using a temperature gradient protocol. dsDNA (non-methylated DNA: lane 2 – 4), ssDNA (non-methylated top strand: lane 5 – 7) and ssDNA (non-methylated bottom strand: lane 8 – 10) were treated with SauUSI either in the presence or absence of ATP and the cleavage pattern was compared with ds/ss DNA controls and run in the presence of ultra low-range DNA ladder.

Interestingly, on treatment of two DNA single strands, which had complimentary sequences, with 100 nM SauUSI, 2 mM ATP and 10 mM magnesium acetate cleavage was observed on a 12 % native-PAGE gel (Figure 1, Lanes 5 - 10). Multiple fragments of ssDNA were produced, which pointed towards the cleavage being sequence-independent (Figure 1, Lane 6 and 9). The efficiency of cleavage seemed to improve in the absence of ATP (Figure 1, Lane 7 and 10).

### ssDNA cleavage: a unique feature of SauUSI, inactivated by nuclease active site mutation

The N-terminal nuclease domain of SauUSI belongs to the Phospholipase-D (PLD) family of protein fold. A characteristic feature of such nucleases is the presence of a strong dimeric interface which brings two ‘HxK’ motifs from each protomer in close proximity to each other to form the enzymatic active site. The histidine side chains from both the protomers act as nucleophiles (29, 30). The histidine in close proximity to the DNA phosphate backbone causes a nucleophilic attack forming a protein-DNA adduct which is resolved by a water molecule whose activation is facilitated by the histidine from the other protomer. This concerted action of two histidines brings about the eventual scission of the phosphate backbone. Mutating the active-site histidine of the nuclease domain to alanine abolished the ssDNA cleavage activity of SauUSI (SauUSI^H119A^) both in the presence and absence of 2 mM ATP (Figure 2a).

**Figure 2:**
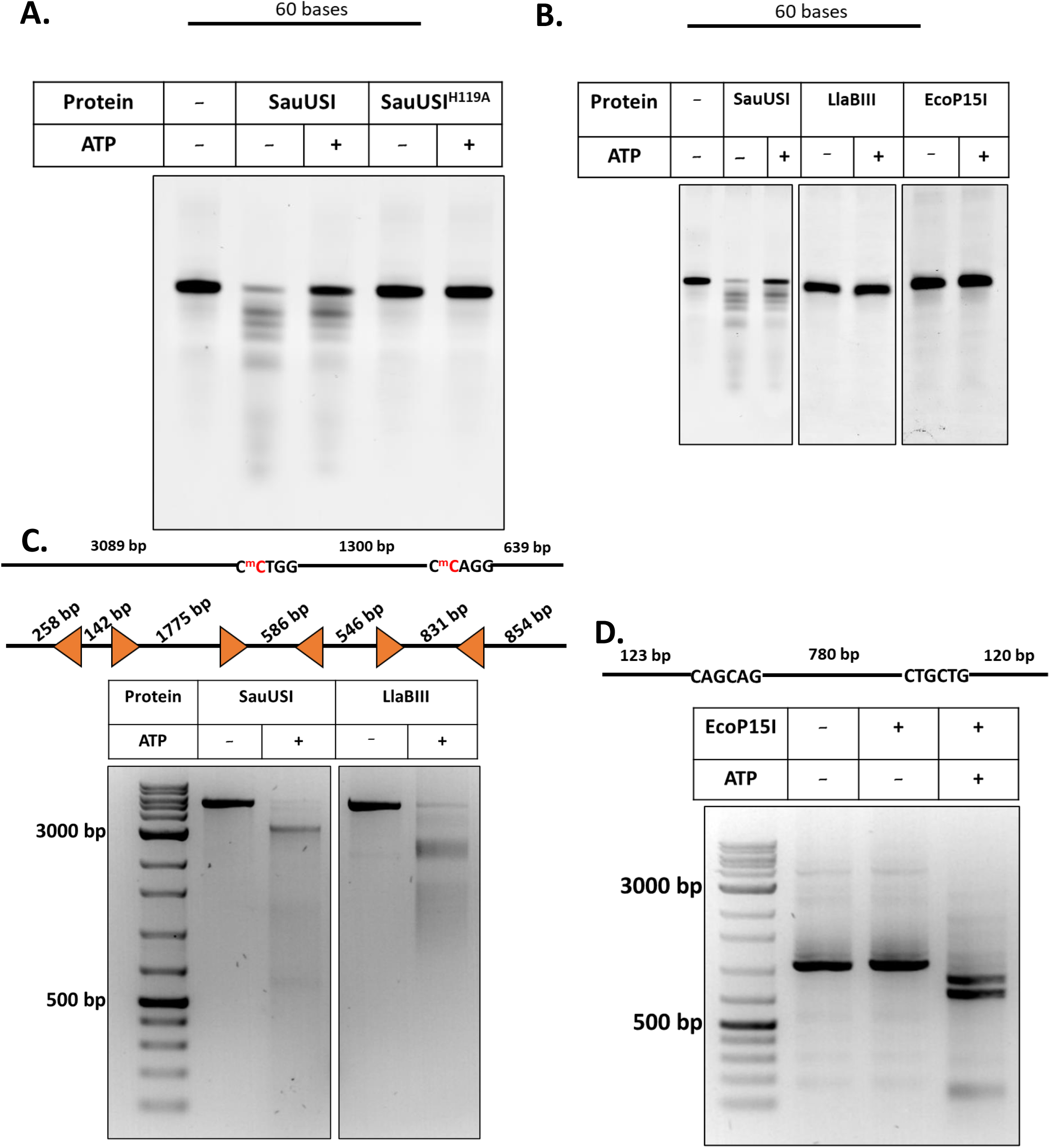
Cleavage of ssDNA is not a feature of all types of ATP-dependent restriction enzymes: (A). Schematic of the substrate used for the cleavage assay. Representative 12 % native PAGE gel to study the cleavage pattern of ssDNA treated with SauUSI and SauUSI H119A (nuclease inactive mutant) either in the presence or absence of ATP. (B) Schematic of the substrate used for the cleavage assay. Representative 12 % native PAGE gel to study the cleavage pattern of ssDNA treated with SauUSI, LlaBIII and EcoP15I either in the presence or absence of ATP. (C) dsDNA substrate used for the cleavage assay. The substrate used was a linearized plasmid and methylation was brought about *in vivo* to generate SauUSI target sites. The dsDNA also had six LlaBIII sites and hence could be used for cleavage. Representative 1 % agarose gel to study the canonical cleavage of SauUSI and LlaBIII in the presence or absence of ATP run in the presence of 1 kb DNAmark ladder (G-Biosciences). (D) dsDNA substrate used for the cleavage assay with EcoP15I. The substrate used was generated *in vitro* by PCR amplification. Representative 1 % agarose gel to study the canonical cleavage of EcoP15I in the presence or absence of ATP.

We tested if other ATP-dependent restriction enzymes LlaBIII (Type ISP RM enzyme) and EcoP15I (Type III RM enzyme) possess the ability to cut ssDNA. The ability of BstNI, a canonical ATP-independent Type II restriction endonuclease that recognizes the same target sequence as SauUSI for dsDNA cleavage, to cleave ssDNA was also tested. Like SauUSI, LlaBIII and EcoP15I require at least two sites in the head-to head orientation for dsDNA cleavage (25, 31–33). 250 nM of ssDNA was treated with 100 nM of LlaBIII and EcoP15I and incubated at 25°C for 1 hour both in the presence and absence of 2 mM ATP (see Materials and Methods for details). Along with these reactions, a control reaction containing 250 nM ssDNA and 100 nM SauUSI either in the presence or absence of 2 mM ATP was set up and incubated at 37°C for 1 hour. Upon performing the aforementioned assay, we concluded that while SauUSI cleaves ssDNA (Figure 2b, Lane 2 and 3). LlaBIII and EcoP15I did not possess this ability (Figure 2b, Lanes 4 -7). BstNI was also not able to cleave ssDNA (Supplementary figure S1). In parallel, we also tested the DNA cleavage activity of these enzymes on their canonical two-site dsDNA under identical conditions. The cleavage of SauUSI, LlaBIII and EcoP15I on their respective substrates gave expected patterns of cleavage in the presence of 2 mM ATP (Figures 2c and 2d).

### ssDNA cleavage does not require divalent cation and is EDTA resistant

Biochemical characterization of the PLD endonuclease Nuc and BfiI revealed that the histidine side chain at the active site functions as a nucleophile thus causing an enzymatic attack of the phosphate backbone of DNA (29, 30). This is in stark contrast to canonical Type II REases which function by coordinating the active-site with two-divalent metal ions thus bringing about the activation of a water molecule which serves as a nucleophile for scission of the phosphate backbone of DNA (34). Therefore, BfiI and Nuc are categorized as metal-independent and EDTA resistant nucleases whereas majority of the Type II REases are metal-dependent and EDTA labile (30, 35). Previous experiments on SauUSI revealed that dsDNA cleavage is stimulated by ATP hydrolysis brought about by the SF2 helicase-like ATPase domain. SF2 helicase-like ATPases require Mg^2+^ ion to bring about stable binding of ATP to the nucleotide binding pocket (36). ATP hydrolysis in SauUSI leads to a confirmational change thereby resulting in translocation along dsDNA.

Since SauUSI possesses a N-terminal PLD nuclease domain whose ssDNA cleavage activity is ATP independent, we tested if it might function as a metal-independent and EDTA resistant nuclease on ssDNA. To this end, we performed a cleavage assay in the presence of the divalent cation chelator EDTA. Since the reaction buffer standardized so far contained 10 mM magnesium acetate, we used a comparable concentration of EDTA (16 mM) to ensure complete chelation of the divalent cations. Upon incubating 200 nM SauUSI with 250 nM ssDNA in 1X NEB4 buffer (cleavage buffer of SauUSI) supplemented with 16 mM EDTA at 37°C for 45 minutes we observed cleavage (Figure 3a. Lane 3 and 4). SauUSI was also able to cleave ssDNA in the absence of magnesium acetate in the reaction mixture (Figure 3a. Lane 5 and 6). Hence, we concluded that ssDNA cleavage by SauUSI does not require divalent cation, which was consistent with the metal-independent nature of other characterized PLD nucleases. We also speculate that many *bona fide* dsDNA endonuclease could also function as ssDNA nuclease that remains to be discovered.

**Figure 3:**
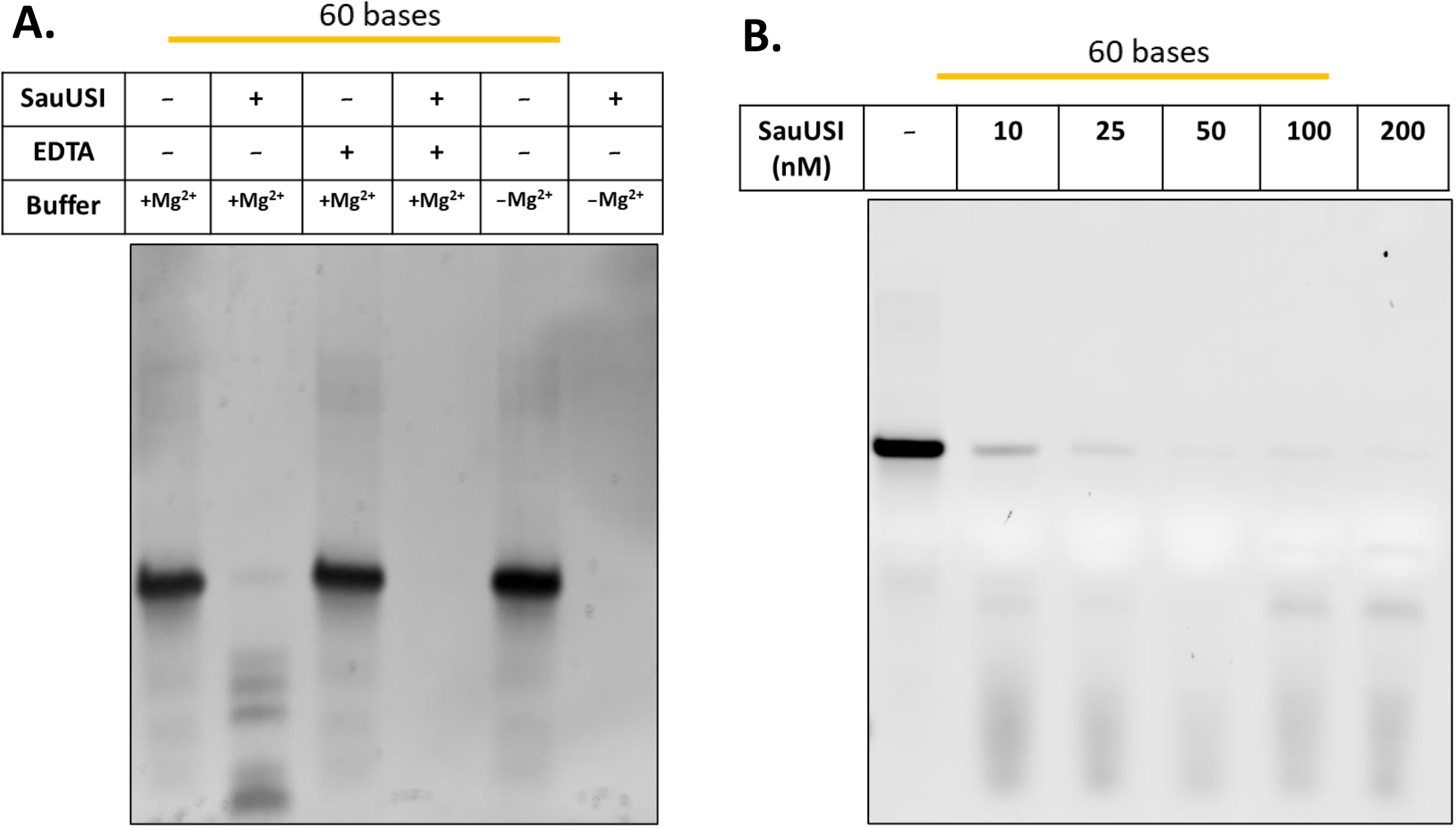
Cleavage of ssDNA does not require divalent cations : (A). Schematic of the ssDNA substrate used for the cleavage assay. Representative 12 % antive PAGE gel to study the cleavage pattern of ssDNA treated with SauUSI either in the presence of EDTA, absence of EDTA or absence of Mg^2+^. (B) Schematic of the ssDNA substrate used for the cleavage assay. Representative 12 % Native PAGE gel to study the cleavage pattern of ssDNA varying the concentration (10 nM to 200 nM) of SauUSI in a reaction buffer free of Mg^2+^.

Interestingly, in the absence of magnesium acetate, SauUSI cleaved ssDNA better than in the presence of it (Figure 3a. Lane 2-6). Further, SauUSI was efficiently able to cleave ssDNA oligos as short as 25 bases in the absence of magnesium acetate (Supplementary figure S2). Performing a cleavage assay with dsDNA possessing two target sites of SauUSI in the absence of 10 mM magnesium acetate showed no detectable cleavage in the presence of ATP whereas, cleavage was achieved on supplementing the reaction buffer with 10 mM magnesium acetate in the presence of ATP (Supplementary figure S3). This was consistent with our assumption that Mg^2+^ was important for ATP hydrolysis by the ATPase domain of SauUSI that activated dsDNA cleavage in a target-site dependent manner. We think that Mg^2+^ is not required for the reaction chemistry of cleavage of the phosphodiester bonds of dsDNA *per se*.

### Single-stranded cleavage is efficient and specific only for DNA and not RNA

Efficient cleavage of 250 nM of ssDNA was observed with as low as 10 nM SauUSI with optimal cleavage noted at 50 nM (Figure 3b and Supplementary figure S4). Time dependent cleavage assay at constant concentration of DNA (250 nM) and protein (50 nM) showed ∼60 % cleavage efficiency within 2 minutes of reaction incubation at 37° C and which increased to ∼90 % by 6 minutes (Figure 4a, 4b). With increasing time, the ssDNA continued to be cut into shorter fragments. This highlighted a potent single-stranded nuclease activity. We also found that the cleavage activity of SauUSI was specific to DNA as it did not cleave ssRNA (Figure 4c). We tested the affinity of SauUSI for ssDNA using electromobility shift assay (EMSA). Varying concentrations of nuclease inactive mutant SauUSI^H119A^ and 250 nM 60-base long ssDNA was used for the experiment. More than 50% of the DNA shifted at a protein concentration of 1.6 μM, which showed that SauUSI had significant affinity for ssDNA (Figure 4d).

**Figure 4:**
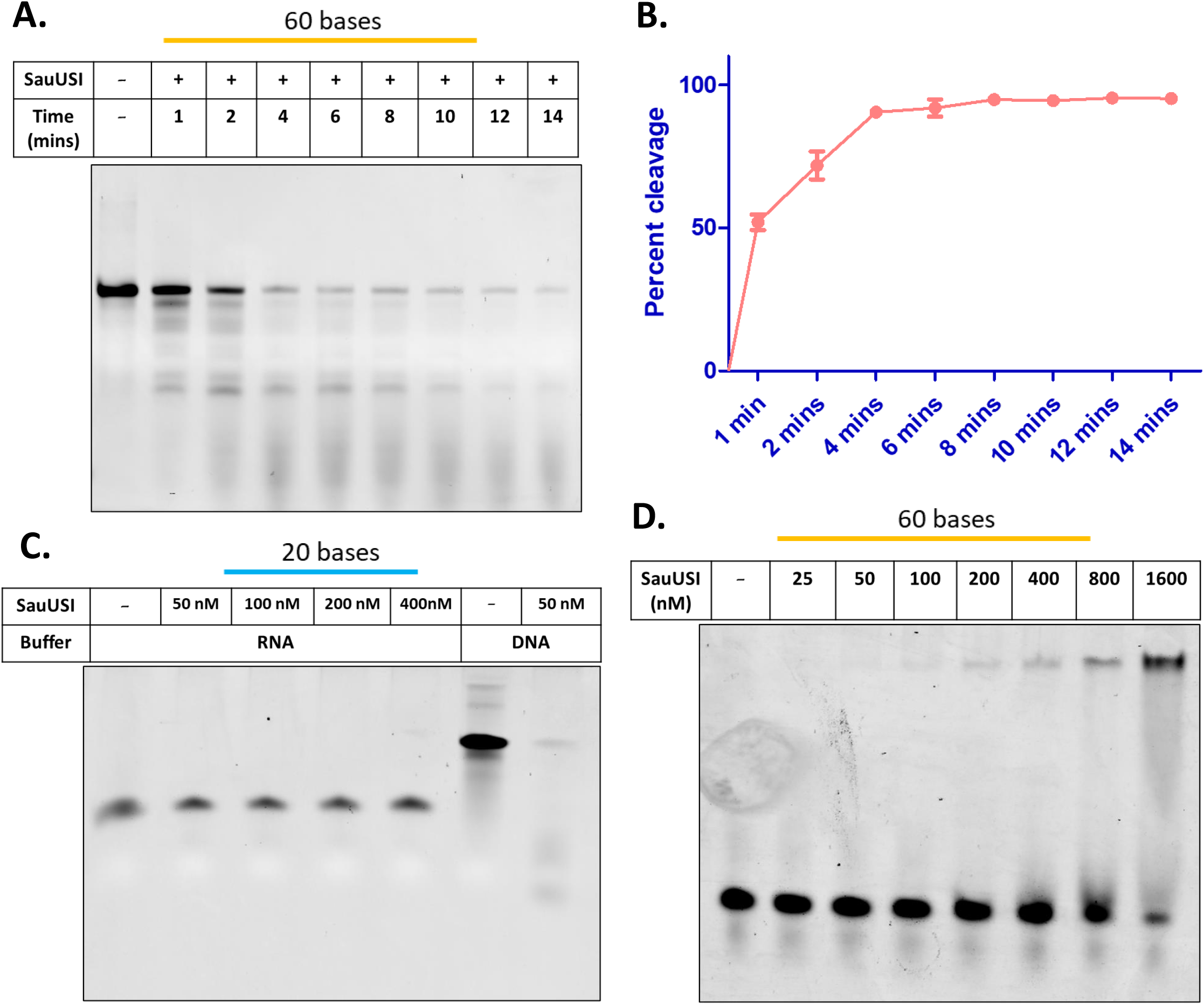
Cleavage of ssDNA is efficient and ssRNA is recalcitrant to cleavage: (A) Schematic of the ssDNA substrate used for the cleavage assay. Representative 12 % native PAGE gel to study the time dependence (1 min – 14 min) of ssDNA cleavage using 50 nM of SauUSI in a reaction buffer free of Mg^2+^. (B) Percent cleavage (y-axis) as a function of time (mins: x-axis) (n=3). (C) Schematic of the ssRNA substrate used for the cleavage assay. Representative 20 % Denaturing PAGE gel to study the cleavage pattern of ssRNA with varying concentration (50 nM 400 nM) of SauUSI. Also shown is the cleavage of a 60-base long ssDNA by 50 nM SauUSI. (D) ssDNA substrate used for EMSA study. Representative 6 % native PAGE to study the binding of ssDNA over a SauUSI concentration range of 25 nM -1600 nM.

### SauUSI is a ssDNA endonuclease

Having established that SauUSI cuts ssDNA efficiently, we proceeded to find if the enzyme is an endonuclease or an exonuclease. Exonucleases are enzymes that trim/cleave DNA from either the 5’ end or 3’ end releasing approximately one nucleotide or a nucleoside at a time (Supplementary figure S5). On the other hand, endonucleases have the ability to cleave DNA within the DNA phosphate backbone without the strict requirement of end trimming (Supplementary figure S5). To distinguish between the two modes of cleavage we used four kinds of substrates: 1) substrates with a 5’ and 3’ overhang, 2) substrate with varying lengths of gaps 3) substrate with a mismatch bubble and 4) substrate with hairpin loop.

Substrates with overhangs were designed to have a 40 bp dsDNA region with either a 20-base long 5’ or 3’ overhang. Cleavage with 50 nM SauUSI did not show any preference for either the 5’ or 3’ termini (Figure 5a). This result indicated that SauUSI did not have a polarity for cleavage and that it might function as an endonuclease. If this were the case then, substrates with a ssDNA gap and mismatch bubble would be labile to cleavage as well. Cleavage assay with a DNA having a 20-base long ssDNA gap flanked by a 15 bp and 25 bp dsDNA region revealed that SauUSI specifically cleaved the ssDNA fragment leaving the dsDNA region unaffected establishing that the enzyme functioned as an endonuclease (Figure 5b. Lanes 1 and 2). Decreasing the ssDNA gap to 15 bases, 10 bases and 5 bases resulted in commensurate reduction in cleavage efficiency (Figure 5b. Lanes 3-8). Furthermore, nicked DNA was not cleaved by SauUSI (Supplementary figure S6). Consistent with these observations, we also found that SauUSI cleaved a DNA having a 15 base mismatch bubble flanked by 22 bp and 23 bp dsDNA. SauUSI selectively cleaved the unannealed single-stranded bubble and not the flanking dsDNA (Figure 5c). As expected, the enzyme also cleaved DNA containing hairpin loop. While cleavage was observed for hairpin having loops of length 20 and 10 bases, a hairpin having a 5-base long loop remained uncleaved (Figure 5d).

**Figure 5:**
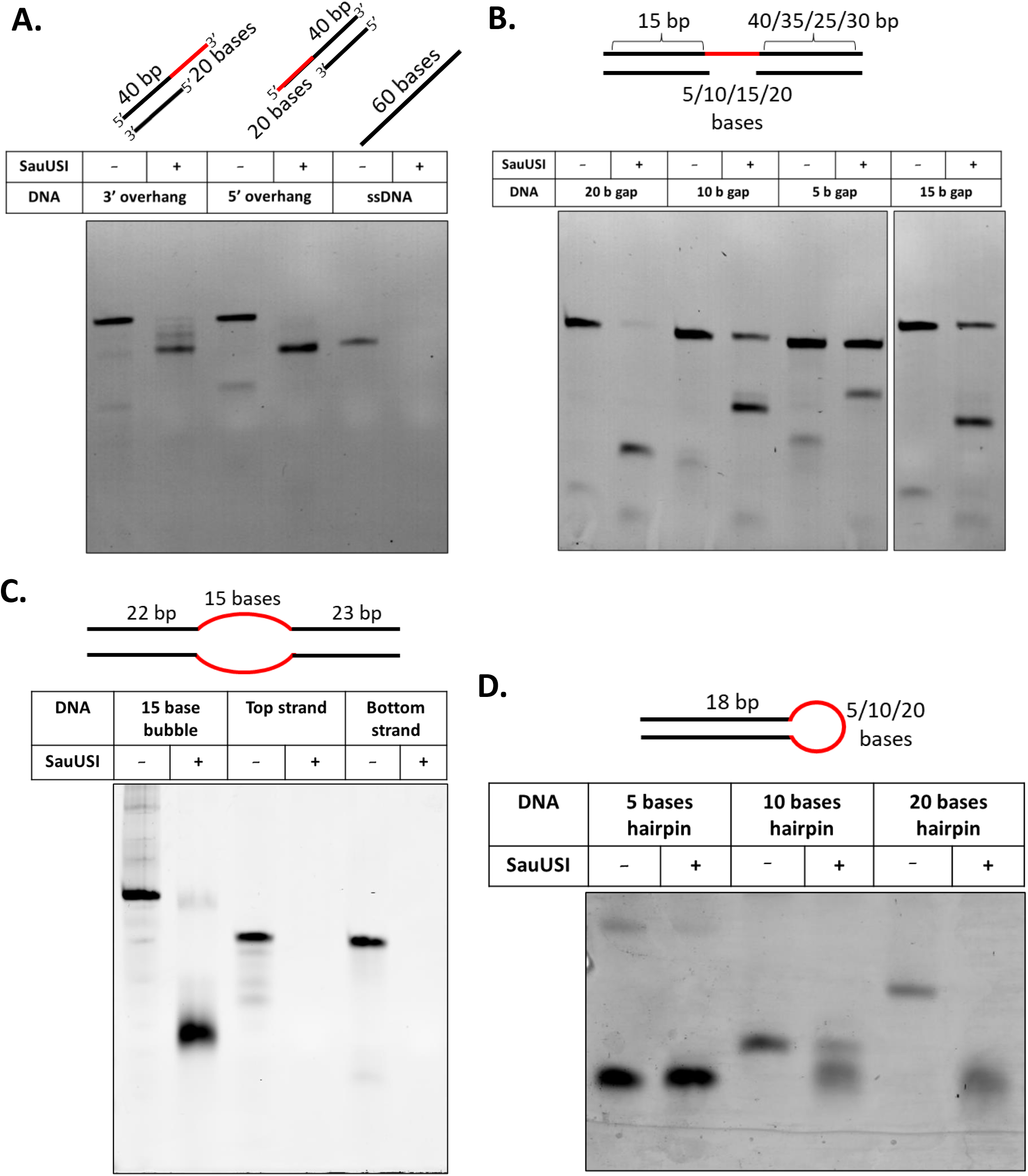
SauUSI is a ssDNA endonuclease: (A). Schematic of the substrates used for the cleavage assay. Representative 12 % native PAGE gel to study the cleavage pattern of overhang substrates and ssDNA treated with SauUSI. (B) Schematic of the DNA substrates with gaps used for the cleavage assay. Representative 12 % native PAGE gel to study the cleavage pattern of DNA substrates with gaps (5/10/15/20 bases gaps) treated with SauUSI. (C) Schematic of the DNA substrate with a mismatch bubble (red) used for the cleavage assay. Representative 12 % Native PAGE gel to study the cleavage pattern of DNA substrate with a mismatch bubble (red) and the individual ssDNA components treated with SauUSI. (D) Schematic of the DNA substrates with a hairpin loop (red) used for the cleavage assay. Representative 12 % native PAGE gel to study the cleavage of DNA hairpins (red: 5/10/20 base hairpin loop) treated with SauUSI.

Structure of SauUSI solved by X-ray crystallography revealed that the nuclease active site of SauUSI is located at the dimeric interface of the two protomers (17). We modelled a ssDNA (adapted from the crystal structure of RecJ, PDB ID: 5F55) along the dimeric interface which matched with our earlier proposal that the path DNA would take to interact with the nuclease active site can be deduced by the position of the three sulfate ions (Figure 6a). The distance between each of the sulfate ions in the crystal structure is ∼12 Å which is almost equivalent to the distance between alternate phosphate moieties in the ssDNA backbone (Supplementary figure S7). Based on the model, it appears that for stable interaction of ssDNA with the SauUSI active site a minimum of 5 phosphate moieties (∼25 Å) would need to interact along the DNA binding path of the nuclease domains (Figure 6b and Supplementary figure S7). The location of the active site residues with respect to the ssDNA indicates that the phosphodiester bond undergoing hydrolysis will be located within the ssDNA polymer rather than at either of the termini in line with our observation that SauUSI is an endonuclease.

**Figure 6:**
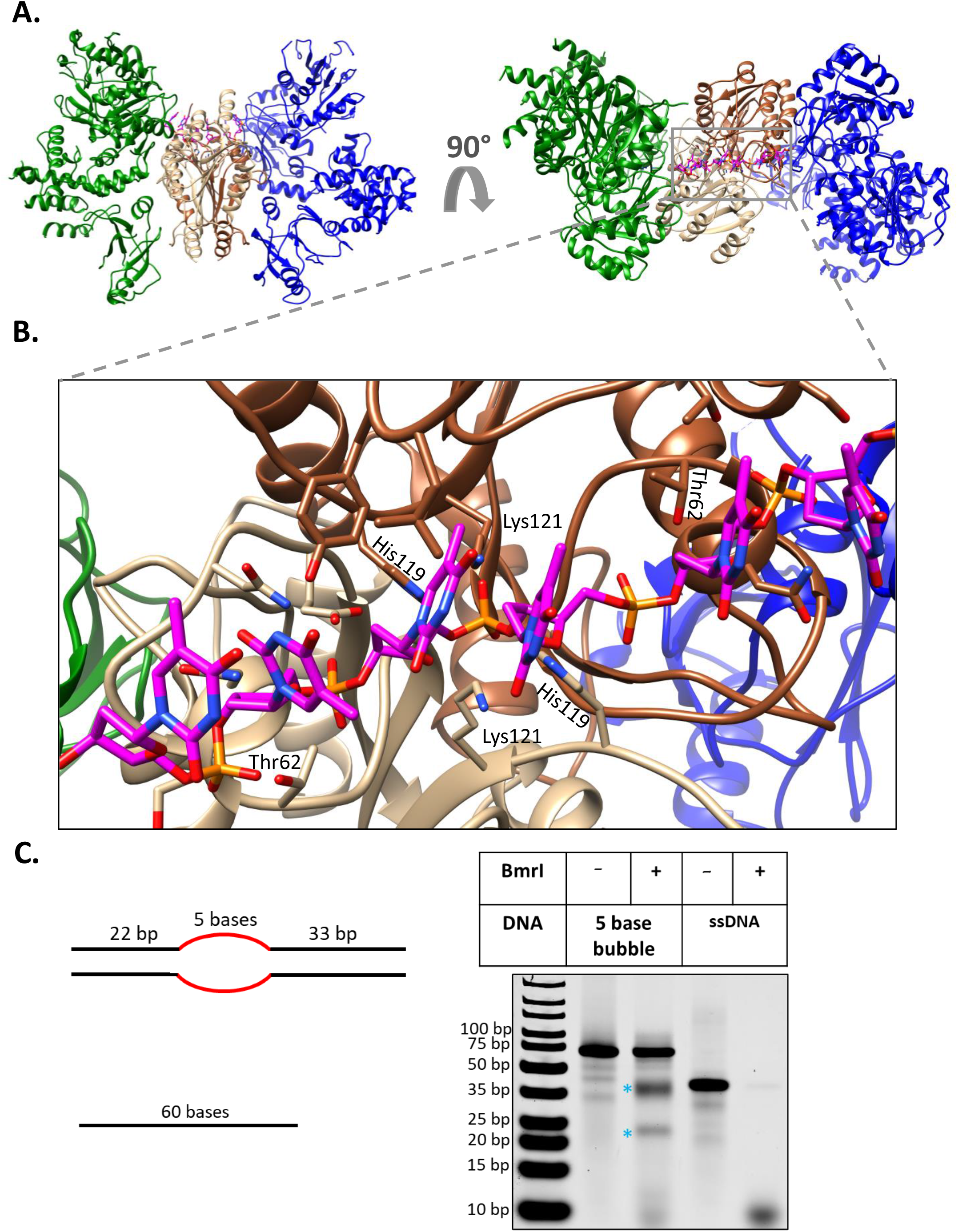
ssDNA modelled along the nuclease active site of SauUSI and ssDNA cleavage by BmrI: (A). Ribbon diagram of two views of the SauUSI crystal structure (PDB: 7CLG) with ssDNA modelled along the nuclease active site. (B) Zoomed in view of the dimeric interface of SauUSI at the two nuclease domains of the protomers interacting with the ssDNA modelled. Some of the active site residues are illustrated. (C) Schematic of the DNA substrate with a mismatch bubble (red) and ssDNA used for the cleavage assay. Representative 12 % Native PAGE gel to study the cleavage pattern of DNA substrate with a mismatch bubble (red) and ssDNA substrates treated with BmrI. Blue astricks represent the flanking dsDNA.

### Cleavage of ssDNA using BmrI

BmrI and BfiI are well characterized nucleotide independent Type IIS REases (29, 35, 37, 38). These enzymes are site-specific DNA cutters which means, they cut at specific locations (5 nucleotides on the top-strand and 4 nucleotides on the bottom strand) close to their respective target sites. Interestingly, these REases can cleave dsDNA even in the absence of divalent cations, a feature that can be attributed to their N-termini which consists of a functional PLD nuclease domain (37–39). Interestingly, it was previously reported that a N-terminal variant of BmrI had the ability to cleave M13mp18 ssDNA (40). We decided to test if BmrI can cleave ssDNA, like the results we obtained for SauUSI. We treated a substrate consisting of a 5 base mismatch bubble flanked by 22 bp and 33 bp dsDNA with BmrI (Figure 6c). Like SauUSI, BmrI selectively cleaved the 5 base mismatch bubble leaving the flanking dsDNA intact (Figure 6c. Lanes 2 and 3) Also, incubating a 60 base-long ssDNA with BmrI we observed efficient cleavage of the ssDNA (Figure 6c. Lanes 4 and 5). These results indicate that ssDNA cleavage could be a feature of REases possessing a PLD nuclease. However, this observation cannot be extended to all other enzymes containing PLD nucleases because there are examples such as Drosophila *melanogaster’s* Zucchini (DmZuc) which is an endoribonuclease involved in piRNA biogenesis that is ineffective against ssDNA substrates (41).

### Factors inhibiting ssDNA cleavage

As mentioned earlier, ssDNA is a transient intermediate of fundamental processes such as replication, transcription and DNA repair. However, ssDNA is sequestered and protected from damage in the cell by single strand DNA binding (SSB) protein. To check if SSB can protect ssDNA from the endonucleolytic activity of SauUSI, we incubated M13mp18 ssDNA with varying concentrations of *E. coli* SSB. We confirmed the binding of SSB to ssDNA based on the shift in the DNA position in an agarose gel on addition of SSB (Figure 7a) (27, 28). Cleavage assay with free M13mp18 ssDNA and SSB-bound M13mp18 ssDNA was carried out. While free M13mp18 ssDNA was nucleolytically degraded, SSB-bound M13mp18 ssDNA was protected from degradation by SauUSI (Figure 7b). This indicated a possible mode of ssDNA protection from SauUSI in the cell.

**Figure 7:**
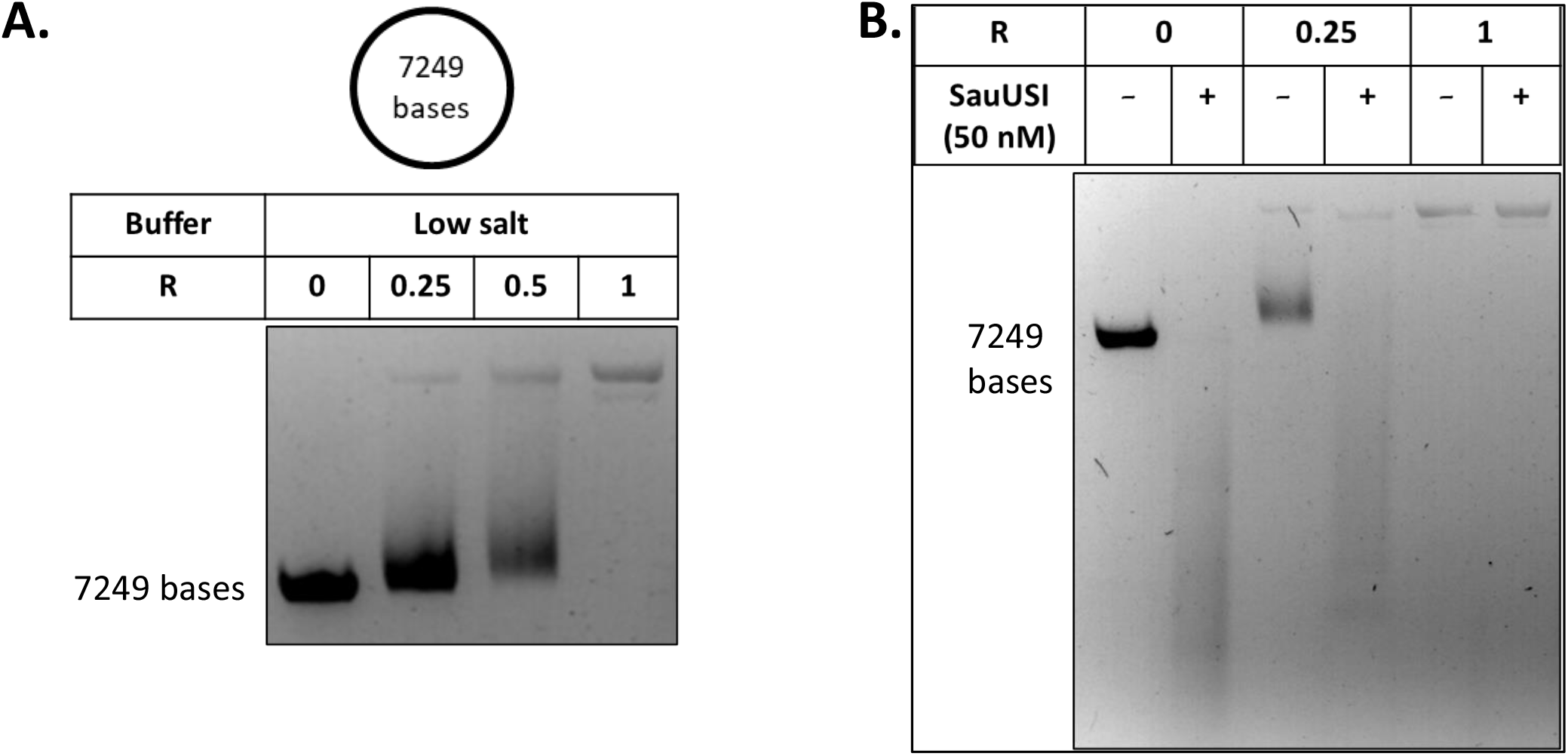
Regulation of ssDNA cleavage by SSB: (A). M13mp18 circular ssDNA used for the cleavage assay. Representative agarose gel to study the binding of *E*.*coli* single-stranded DNA binding (SSB) protein at different R values (ratio of SSB:ssDNA). (B) Representative agarose gel to study the cleavage pattern of unbound or bound M13mp18 (bound to *E*.*coli* SSB) treated with SauUSI performed over R values 0, 0.25 and 1.

We had earlier found that the presence of Mg^2+^ lowered the ssDNA nuclease activity of SauUSI (Figure 3a). We had also noticed that ATP lowered the ssDNA cleavage activity (Figures 1, 2a and 2b). We studied this effect further by performing the cleavage assay by varying the concentrations of ATP from 50 μM to 4 mM. A reduction in efficiency of cleavage was observed as the concentration of ATP increased (Figure 8a). The inhibition was >85 % at 4 mM ATP and the IC_50_ calculated was ∼ 760 μM (Figure 8b). We next tested if other nucleoside tri-phosphates possess the ability to inhibit cleavage. ATP and dATP conferred maximum inhibition to cleavage, whereas inhibition by GTP, CTP and UTP was comparatively lower (Figure 8c). ADP, on the other hand, appeared as inhibitory as ATP, while AMP had limited inhibition (Figure 8d). We gathered further evidence for the inhibitory effect of ATP and Mg^2+^ by performing a cleavage assay on a 200-base long and 36-base long ssDNA molecules (Figure 8e). As expected, when we incubated either 200-base long or 36-base long ssDNA with SauUSI in the absence of Mg^2+^ and ATP (Figure 8e, Lane 2 and 5) there was efficient cleavage. However, in the presence of 10 mM Mg^2+^ and 4 mM ATP (Figure 8e, Lane 3 and 6) the cleavage efficiency of SauUSI drastically reduced. This result suggested a regulatory role for the ATPase domain in the ssDNA cleavage activity.

**Figure 8:**
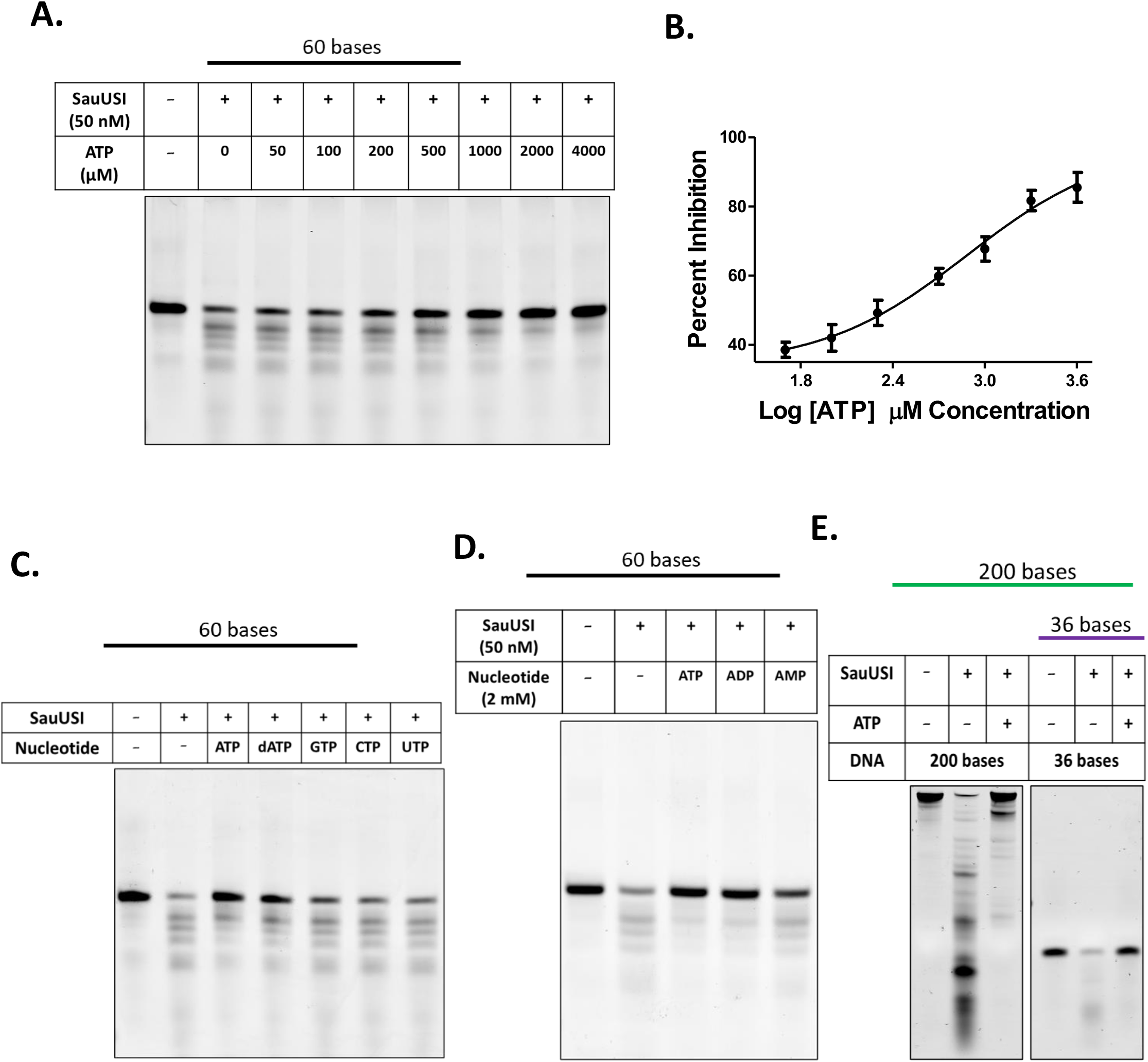
Regulation of ssDNA cleavage by nucleotides: (A) Schematic of the ssDNA substrate used for the cleavage assay. Representative 12 % native PAGE gel to study the effect of an increasing concentration (0 – 4000 μ M) of ATP on the cleavage of ssDNA by SauUSI. (B) Percent inhibition plotted as a function of Log ATP concentration to calculate IC_50_ value for the inhibition (C) Schematic of the ssDNA substrate used for the cleavage assay. Representative 12 % native PAGE gel to study the effect of different nucleoside triphosphates on the cleavage of ssDNA by SauUSI. (D) Schematic of the ssDNA substrate used for the cleavage assay. Representative 12 % native PAGE gel to study the effect of different adenine nucleotides (ATP, ADP and AMP) on the cleavage of ssDNA by SauUSI. (E) Schematic of the ssDNA used for the cleavage assay. Representative 12 % native PAGE gel to study the inhibitory effect of ATP (4 mM) on cleavage of a 200 base and 36 bases ssDNA by SauUSI

## Discussion

We have demonstrated here that the restriction enzyme SauUSI has the ability to endonucleolytically cleave ssDNA. This is, to the best of our knowledge, the first example of a restriction enzyme being a *bona fide* ssDNA endonuclease. While there have been previous reports of restriction enzymes cleaving ssDNA, the cleavage was concluded to be the result of formation of transient dsDNA regions that were susceptible to cleavage. The observations supporting this conclusion were (i) the requirement of the same target site for ssDNA cleavage as for dsDNA cleavage (11, 13); (ii) correlation between the stability of the dsDNA structure and efficiency of cleavage (11, 16); the requirement of higher enzyme concentration for ssDNA cleavage than dsDNA cleavage (8–15) possibly due to the transient nature of the dsDNA substrate (16). In contrast, cleavage of ssDNA by SauUSI did not show any particular sequence requirement. Also, efficient ssDNA cleavage required only as much concentration of the enzyme as dsDNA cleavage; and did not require ATP – a requisite for dsDNA cleavage by SauUSI.

*S. aureus* is a notorious pathogen known to cause various human maladies ranging from minor skin infections to life threatening endocarditis (42, 43). The success of the organism as a pathogen is greatly determined by its ability to evade host immune response and acquire antibiotic resistance (44–48). Micrococcal Nuclease (MN), a secreted endonuclease encoded by *nuc1*, and the cell-wall bound nuclease encoded by *nuc2* are involved in immune evasion thus increasing pathogenesis (49–52). The diverse Cas nuclease repertoire of the CRISPR systems and the different types of R-M systems in *S. aureus* regulate the acquisition of foreign genetic material (2, 18, 53–56). The dsDNA cleavage activity of SauUSI has been identified as a potent barrier to the entry of methylated dsDNA either by transduction or transformation (2, 17–19). However, we also find that SauUSI can also function as a potent ssDNA endonuclease. Consequently, SauUSI can be a barrier to invasion of ssDNA phages or to the process of conjugation, which involves receipt of chromosomal or extrachromosomal material as ssDNA from donor bacteria (57). Conjugation between the donor *Enterococcus sp*. and the recipient *S. aureus* is a major mode of acquisition of antibiotic resistance (20).

The study also demonstrated that the enzyme endonucleolytically cleaved ssDNA in a variety of DNA structures, including DNA with single-stranded overhangs, gaps, bubbles and loops. Efficiency of cleavage of ssDNA gaps decreased with decreasing length of the gap and SauUSI did not cut nicked DNA. These ssDNA containing structures occur transiently but frequently in a cell at various stages of its physiology, including during DNA replication, repair and recombination. Consequently, these important DNA structural intermediates will be susceptible to the ssDNA cleavage activity of SauUSI, if left unregulated. We found that the single-stranded endonucleolytic activity of SauUSI was inhibited by Mg^2+^, ATP, dATP or ADP. However, inhibitory effects of AMP were much lower. The IC_50_ of ATP inhibition was measured to be 760 μM, a concentration which is lower than the concentrations of ATP in bacteria under normal physiological conditions (58).

Inhibition of the ssDNA nuclease activity of SauUSI by either ATP or ADP indicates that the inhibition is due to nucleotide binding rather than hydrolysis. Inhibition of the nuclease could be a result of conformational changes in SauUSI resulting from nucleotide binding. A similar proposal has been made for the regulation of ATPase-coupled nuclease *S. aureus* DinG (sarDinG) by ATP (59). sarDinG has a nuclease domain and an inactive helicase that can bind and hydrolyze ATP but cannot unwind DNA (59). The nuclease has a ssDNA 3’-exonuclease activity, which is negatively regulated by ATP concentration (59). It is proposed that ATP binding at the helicase domain causes a conformational change that inhibits the exonuclease. Apart from the inhibitory effect of ATP/ADP and Mg^2+^ on ssDNA cleavage activity of SauUSI, we found that SSB also prevented this cleavage *in vitro*. Thenceforth, we think that the ssDNA cleavage activity of SauUSI is tightly regulated under physiological conditions. This also leads us to question if the ssDNA cleavage activity of SauUSI would ever manifest and serve as a bacterial defense against foreign ssDNA.

We speculate that SauUSI could function as a bacterial defense against single-stranded DNA phages in a manner analogous to the nicking endonuclease GajA of the nucleotide sensing Gabija system in *Bacillus sp* (60). Like SauUSI, GajA is negatively regulated by nucleotide concentrations (dNTPs, NTPs, dNDPs and NDPs), while monophosphate nucleotides (ribo and deoxyribo derivatives) and nucleosides do not impede cleavage (60). It is proposed that during phage infection, active DNA transcription and replication deplete NTP and dNTP pool. Decrease in the trinucleotide pool would then activate the GajA nuclease causing degradation of phage DNA and host genome, resulting in abortive infection (60). A similar scenario can pan out when bacteria hosting SauUSI are infected by single-stranded DNA phages. Depletion of NTPs and dNTPs during phage DNA replication and transcription will activate cleavage of ssDNA, including the replicating phage genome and that formed transiently in the host genome, causing abortive infection. If the foreign DNA entering the host bacteria is double-stranded and has the SauUSI target sites, shredding of the foreign dsDNA would be the prominent barrier to the entry (17).

*S. aureus* is a facultative intracellular pathogen and, hence, is subject to a host of DNA damaging agents such as reactive oxygen species (ROS) synthesized by the host defense mechanisms (61). As a result, they have evolved numerous DNA repair mechanisms that involves multiple nucleases acting in tandem (62). *sbcC* and *sbcD* encode the endonuclease SbcCD to resolve stalled replication forks (63); a homologue of YqfS is known to be involved in base-excision repair (62); UvrABC endonuclease complex plays a pivotal role in nucleotide-excision repair (62). The mismatch repair system, which helps correct aberrant incorporation of nucleotides during replication, is thought to be brought about by the nicking-endonuclease MutL (because of the absence of homologs of MutH) and excision of the erroneous region by the exonuclease RecJ (62, 64). dsDNA breaks are repaired by homologous recombination in *S. aureus*, which is initiated by trimming the ends by the RexAB helicase-nuclease to form 3’-overhangs crucial for the process of strand invasion (65). The holiday junction so formed is further resolved by another nuclease MutS2 (62, 66). SauUSI having the ability to cut a mismatch bubble, a ssDNA gap or an overhang, has the potential to moonlights as a DNA repair endonuclease *in-vivo*.

The ability of SauUSI to endonucleolytically cleave ssDNA leads to a variety of interesting possibilities and roles for the enzyme *in vivo* that remains to be unraveled. However, the ability of an enzyme to carry out a specific reaction does not necessarily imply an *in vivo* significance for that enzymatic reaction. An example of such a scenario is that noted for CRISPR-Cas12a recently. CRISPR-Cas12 has a strong ssDNA endonucleolytic activity *in vitro* (67), which does not have a physiological relevance (68). The present study provides a platform to perform detailed studies to understand the physiological role of the ssDNA cleavage activity of SauUSI, in particular during the process of transduction and conjugation.

## Acknowledgements

We thank IISER Pune for the laboratory infrastructure and the common equipment facility. We thank Dr. Om Prakash Chouhan, Akhilesh Pratap Singh and Mahesh Kumar Chand for providing purified EcoP15I and LlaBIII. We thank Singh Akash Kailashprasad for performing some cleavage assays during his training period in the laboratory. We also thank other lab members for their constructive inputs during experimentation and preparation of the manuscript.

## References

1. Wheatley, R.M. and MacLean, R.C. (2021) CRISPR-Cas systems restrict horizontal gene transfer in Pseudomonas aeruginosa. ISME J, 15, 1420–1433.

2. Corvaglia, A.R., François, P., Hernandez, D., Perron, K., Linder, P. and Schrenzel, J. (2010) A type III-like restriction endonuclease functions as a major barrier to horizontal gene transfer in clinical Staphylococcus aureus strains. Proc Natl Acad Sci U S A, 107, 11954–11958.

3. Price, V.J., McBride, S.W., Hullahalli, K., Chatterjee, A., Duerkop, B.A. and Palmer, K.L. (2019) Enterococcus faecalis CRISPR-Cas Is a Robust Barrier to Conjugative Antibiotic Resistance Dissemination in the Murine Intestine. mSphere, 4.

4. Wang, G., Song, G. and Xu, Y. (2020) Association of CRISPR/Cas System with the Drug Resistance in Klebsiella pneumoniae. Infect Drug Resist, 13, 1929–1935.

5. Tyumentseva, M., Mikhaylova, Y., Prelovskaya, A., Tyumentsev, A., Petrova, L., Fomina, V., Zamyatin, M., Shelenkov, A. and Akimkin, V. (2021) Genomic and Phenotypic Analysis of Multidrug-Resistant Acinetobacter baumannii Clinical Isolates Carrying Different Types of CRISPR/Cas Systems. Pathogens, 10.

6. Nishino, T. and Morikawa, K. (2002) Structure and function of nucleases in DNA repair: shape, grip and blade of the DNA scissors. Oncogene, 21, 9022–9032.

7. Bernheim, A. and Sorek, R. (2020) The pan-immune system of bacteria: antiviral defence as a community resource. Nat Rev Microbiol, 18, 113–119.

8. Blakesley, R.W. and Wells, R.D. (1975) ‘Single-stranded’ DNA from phiX174 and M13 is cleaved by certain restriction endonucleases. Nature, 257, 421–422.

9. Horiuchi, K. and Zinder, N.D. (1975) Site-Specific Cleavage of Single-Stranded DNA by a Hemophilus Restriction Endonuclease. Proc Natl Acad Sci U S A, 72, 2555–2558.

10. Reckmann, B. and Krauss, G. (1987) The cleavage of single-stranded DNA by the isoschizomeric restriction endonuclease HhaI and CfoI. Biochim Biophys Acta, 908, 90–96.

11. Nishigaki, K., Kaneko, Y., Wakuda, H., Husimi, Y. and Tanaka, T. (1985) Type II restriction endonucleases cleave single-stranded DNAs in general. Nucleic Acids Res, 13, 5747–5760.

12. Blakesley, R.W., Dodgson, J.B., Nes, I.F. and Wells, R.D. (1977) Duplex regions in ‘single-stranded’ phiX174 DNA are cleaved by a restriction endonuclease from Haemophilus aegyptius. J Biol Chem, 252, 7300–7306.

13. Godson, G.N. and Roberts, R.J. (1976) dna, single stranded/*metab. Virology, 73, 561–567.

14. Yoo, O.J. and Agarwal, K.L. (1980) Cleavage of single strand oligonucleotides and bacteriophage phi X174 DNA by Msp I endonuclease. J Biol Chem, 255 22, 10559–10562.

15. Hofer, B., Ruhe, G., Koch, A. and Köster, H. (1982) Primary and secondary structure specificity of the cleavage of ‘single-stranded’ DNA by endonuclease Hinf I. Nucleic Acids Res, 10, 2763–2773.

16. Nishigaki, K. (2018) Short lifetime structures appearing in RNA and DNA. Brief Funct Genomics, 18, 174–181.

17. Tumuluri, V.S., Rajgor, V., Xu, S.-Y., Chouhan, O.P. and Saikrishnan, K. (2021) Mechanism of DNA cleavage by the endonuclease SauUSI: a major barrier to horizontal gene transfer and antibiotic resistance in Staphylococcus aureus. Nucleic Acids Res, 49, 2161–2178.

18. Xu, S.-Y., Corvaglia, A.R., Chan, S.-H., Zheng, Y. and Linder, P. (2011) A type IV modification-dependent restriction enzyme SauUSI from Staphylococcus aureus subsp. aureus USA300. Nucleic Acids Res, 39, 5597–5610.

19. Monk, I.R., Shah, I.M., Xu, M., Tan, M.-W. and Foster, T.J. (2012) Transforming the untransformable: application of direct transformation to manipulate genetically Staphylococcus aureus and Staphylococcus epidermidis. mBio, 3.

20. de Niederhäusern, S., Bondi, M., Messi, P., Iseppi, R., Sabia, C., Manicardi, G. and Anacarso, I. (2011) Vancomycin-resistance Transferability from VanA Enterococci to Staphylococcus aureus. Curr Microbiol, 62, 1363–1367.

21. Noble, W.C., Virani, Z. and Cree, R.G. (1992) Co-transfer of vancomycin and other resistance genes from Enterococcus faecalis NCTC 12201 to Staphylococcus aureus. FEMS Microbiol Lett, 72, 195–198.

22. Périchon, B. and Courvalin, P. (2009) VanA-type vancomycin-resistant Staphylococcus aureus. Antimicrob Agents Chemother, 53, 4580–4587.

23. Maree, M., Thi Nguyen, L.T., Ohniwa, R.L., Higashide, M., Msadek, T. and Morikawa, K. (2022) Natural transformation allows transfer of SCCmec-mediated methicillin resistance in Staphylococcus aureus biofilms. Nat Commun, 13, 2477.

24. Tumuluri, V.S. and Saikrishnan, K. (2022) Heterologous Expression and High Degree Purification of the Restriction Endonuclease SauUSI. Bio Protoc, 12, e4275.

25. Chand, M.K., Nirwan, N., Diffin, F.M., van Aelst, K., Kulkarni, M., Pernstich, C., Szczelkun, M.D. and Saikrishnan, K. (2015) Translocation-coupled DNA cleavage by the Type ISP restriction-modification enzymes. Nat Chem Biol, 11, 870–877.

26. Ahmad, I., Kulkarni, M., Gopinath, A. and Saikrishnan, K. (2018) Single-site DNA cleavage by Type III restriction endonuclease requires a site-bound enzyme and a trans-acting enzyme that are ATPase-activated. Nucleic Acids Res, 46, 6229–6237.

27. Lohman, T.M., Overman, L.B. and Datta, S. (1986) Salt-dependent changes in the DNA binding co-operativity of Escherichia coli single strand binding protein. J Mol Biol, 187, 603–615.

28. Waldman, V.M., Weiland, E., Kozlov, A.G. and Lohman, T.M. (2016) Is a fully wrapped SSB-DNA complex essential for Escherichia coli survival? Nucleic Acids Res, 44, 4317–4329.

29. Sasnauskas, G., Halford, S.E. and Siksnys, V. (2003) How the BfiI restriction enzyme uses one active site to cut two DNA strands. Proc Natl Acad Sci U S A, 100, 6410–6415.

30. Gottlin, E.B., Rudolph, A.E., Zhao, Y., Matthews, H.R. and Dixon, J.E. (1998) Catalytic mechanism of the phospholipase D superfamily proceeds via a covalent phosphohistidine intermediate. Proc Natl Acad Sci U S A, 95, 9202–9207.

31. Ramanathan, S.P., van Aelst, K., Sears, A., Peakman, L.J., Diffin, F.M., Szczelkun, M.D. and Seidel, R. (2009) Type III restriction enzymes communicate in 1D without looping between their target sites. Proc Natl Acad Sci U S A, 106, 1748–1753.

32. van Aelst, K., Tóth, J., Ramanathan, S.P., Schwarz, F.W., Seidel, R. and Szczelkun, M.D. (2010) Type III restriction enzymes cleave DNA by long-range interaction between sites in both head-to-head and tail-to-tail inverted repeat. Proc Natl Acad Sci U S A, 107, 9123–9128.

33. Dryden, D.T., Murray, N.E. and Rao, D.N. (2001) Nucleoside triphosphate-dependent restriction enzymes. Nucleic Acids Res, 29, 3728–3741.

34. Pingoud, A., Fuxreiter, M., Pingoud, V. and Wende, W. (2005) Type II restriction endonucleases: structure and mechanism. Cell Mol Life Sci, 62, 685–707.

35. Zaremba, M., Urbanke, C., Halford, S.E. and Siksnys, V. (2004) Generation of the BfiI Restriction Endonuclease from the Fusion of a DNA Recognition Domain to a Non-specific Nuclease from the Phospholipase D Superfamily. J Mol Biol, 336, 81–92.

36. Fairman-Williams, M.E., Guenther, U.-P. and Jankowsky, E. (2010) SF1 and SF2 helicases: family matters. Curr Opin Struct Biol, 20, 313–324.

37. Chan, S., Bao, Y., Ciszak, E., Laget, S. and Xu, S. (2007) Catalytic domain of restriction endonuclease BmrI as a cleavage module for engineering endonucleases with novel substrate specificities. Nucleic Acids Res, 35, 6238–6248.

38. Bao, Y., Higgins, L., Zhang, P., Chan, S., Laget, S., Sweeney, S., Lunnen, K. and Xu, S. (2008) Expression and purification of BmrI restriction endonuclease and its N-terminal cleavage domain variants. Protein Expr Purif, 58, 42–52.

39. Sapranauskas, R., Sasnauskas, G., Lagunavicius, A., Vilkaitis, G., Lubys, A. and Siksnys, V. (2000) Novel Subtype of Type IIs Restriction Enzymes: BfiI endonuclease exhibits similarities to the EDTA-resistant nuclease Nuc of Salmonella typhimurium. Journal of Biological Chemistry, 275, 30878–30885.

40. Chan, S., Bao, Y., Ciszak, E., Laget, S. and Xu, S. (2007) Catalytic domain of restriction endonuclease BmrI as a cleavage module for engineering endonucleases with novel substrate specificities. Nucleic Acids Res, 35, 6238–6248.

41. Nishimasu, H., Ishizu, H., Saito, K., Fukuhara, S., Kamatani, M.K., Bonnefond, L., Matsumoto, N., Nishizawa, T., Nakanaga, K., Aoki, J., et al. (2012) Structure and function of Zucchini endoribonuclease in piRNA biogenesis. Nature, 491, 284–287.

42. Tong, S.Y.C., Davis, J.S., Eichenberger, E., Holland, T.L. and Fowler, V.G.J. (2015) Staphylococcus aureus infections: epidemiology, pathophysiology, clinical manifestations, and management. Clin Microbiol Rev, 28, 603–661.

43. David, M.Z. and Daum, R.S. (2010) Community-Associated Methicillin-Resistant Staphylococcus aureus: Epidemiology and Clinical Consequences of an Emerging Epidemic. Clin Microbiol Rev, 23, 616–687.

44. Foster, T.J., Geoghegan, J.A., Ganesh, V.K. and Höök, M. (2014) Adhesion, invasion and evasion: the many functions of the surface proteins of Staphylococcus aureus. Nat Rev Microbiol, 12, 49–62.

45. Itoh, S., Hamada, E., Kamoshida, G., Yokoyama, R., Takii, T., Onozaki, K. and Tsuji, T. (2010) Staphylococcal superantigen-like protein 10 (SSL10) binds to human immunoglobulin G (IgG) and inhibits complement activation via the classical pathway. Mol Immunol, 47, 932–938.

46. Pietrocola, G., Nobile, G., Rindi, S. and Speziale, P. (2017) Staphylococcus aureus Manipulates Innate Immunity through Own and Host-Expressed Proteases. Front Cell Infect Microbiol, 7.

47. Pantosti, A., Sanchini, A. and Monaco, M. (2007) Mechanisms of antibiotic resistance in Staphylococcus aureus. Future Microbiol, 2, 323–334.

48. Ito, T., Okuma, K., Ma, X.X., Yuzawa, H. and Hiramatsu, K. (2003) Insights on antibiotic resistance of Staphylococcus aureus from its whole genome: genomic island SCC. Drug Resist Updat, 6, 41–52.

49. Tang, J., Zhou, R., Shi, X., Kang, M., Wang, H. and Chen, H. (2008) Two thermostable nucleases coexisted in Staphylococcus aureus: evidence from mutagenesis and in vitro expression. FEMS Microbiol Lett, 284, 176–183.

50. Thammavongsa, V., Missiakas, D.M. and Schneewind, O. (2013) Staphylococcus aureus Degrades Neutrophil Extracellular Traps to Promote Immune Cell Death. Science (1979), 342, 863–866.

51. Kiedrowski, M.R., Kavanaugh, J.S., Malone, C.L., Mootz, J.M., Voyich, J.M., Smeltzer, M.S., Bayles, K.W. and Horswill, A.R. (2011) Nuclease modulates biofilm formation in community-associated methicillin-resistant Staphylococcus aureus. PLoS One, 6, e26714.

52. Yu, J., Jiang, F., Zhang, F., Hamushan, M., Du, J., Mao, Y., Wang, Q., Han, P., Tang, J. and Shen, H. (2021) Thermonucleases Contribute to Staphylococcus aureus Biofilm Formation in Implant-Associated Infections-A Redundant and Complementary Story. Front Microbiol, 12, 687888.

53. Sussenbach, J.S., Monfoort, C.H., Schiphof, R. and Stobberingh, E.E. (1976) A restriction endonuclease from Staphylococcus aureus. Nucleic Acids Res, 3, 3193–3202.

54. Waldron, D.E. and Lindsay, J.A. (2006) Sau1: a novel lineage-specific type I restriction-modification system that blocks horizontal gene transfer into Staphylococcus aureus and between S. aureus isolates of different lineages. J Bacteriol, 188, 5578–5585.

55. Cruz-López, E.A., Rivera, G., Cruz-Hernández, M.A., Martínez-Vázquez, A.V., Castro-Escarpulli, G., Flores-Magallón, R., Vázquez, K., Cruz-Pulido, W.L. and Bocanegra-García, V. (2021) Identification and Characterization of the CRISPR/Cas System in Staphylococcus aureus Strains From Diverse Sources. Front Microbiol, 12, 656996.

56. Nishimasu, H., Cong, L., Yan, W.X., Ran, F.A., Zetsche, B., Li, Y., Kurabayashi, A., Ishitani, R., Zhang, F. and Nureki, O. (2015) Crystal Structure of Staphylococcus aureus Cas9. Cell, 162, 1113–1126.

57. Olsen, N.S., Forero-Junco, L., Kot, W. and Hansen, L.H. (2020) Exploring the Remarkable Diversity of Culturable Escherichia coli Phages in the Danish Wastewater Environment. Viruses, 12.

58. Buckstein, M.H., He, J. and Rubin, H. (2008) Characterization of nucleotide pools as a function of physiological state in Escherichia coli. J Bacteriol, 190, 718–726.

59. McRobbie, A.-M., Meyer, B., Rouillon, C., Petrovic-Stojanovska, B., Liu, H. and White, M.F. (2012) Staphylococcus aureus DinG, a helicase that has evolved into a nuclease. Biochem J, 442, 77–84.

60. Cheng, R., Huang, F., Wu, H., Lu, X., Yan, Y., Yu, B., Wang, X. and Zhu, B. (2021) A nucleotide-sensing endonuclease from the Gabija bacterial defense system. Nucleic Acids Res, 49, 5216–5229.

61. Ha, K.P., Clarke, R.S., Kim, G.-L., Brittan, J.L., Rowley, J.E., Mavridou, D.A.I., Parker, D., Clarke, T.B., Nobbs, A.H. and Edwards, A.M. (2020) Staphylococcal DNA Repair Is Required for Infection. mBio, 11.

62. Ha, K.P. and Edwards, A.M. (2021) DNA Repair in Staphylococcus aureus. Microbiol Mol Biol Rev, 85, e0009121.

63. Cirz, R.T., Jones, M.B., Gingles, N.A., Minogue, T.D., Jarrahi, B., Peterson, S.N. and Romesberg, F.E. (2007) Complete and SOS-Mediated Response of Staphylococcus aureus to the Antibiotic Ciprofloxacin. J Bacteriol, 189, 531–539.

64. Prunier, A.-L. and Leclercq, R. (2005) Role of mutS and mutL genes in hypermutability and recombination in Staphylococcus aureus. J Bacteriol, 187, 3455–3464.

65. Clarke, R.S., Ha, K.P. and Edwards, A.M. (2021) RexAB Promotes the Survival of Staphylococcus aureus Exposed to Multiple Classes of Antibiotics. Antimicrob Agents Chemother, 65, e0059421.

66. Burby, P.E. and Simmons, L.A. (2017) MutS2 Promotes Homologous Recombination in Bacillus subtilis. J Bacteriol, 199.

67. Chen, J.S., Ma, E., Harrington, L.B. Costa, M. Da, Tian, X., Palefsky, J.M. and Doudna, J.A. (2018) CRISPR-Cas12a target binding unleashes indiscriminate single-stranded DNase activity. Science (1979), 360, 436–439.

68. Marino, N.D., Pinilla-Redondo, R. and Bondy-Denomy, J. (2022) CRISPR-Cas12a targeting of ssDNA plays no detectable role in immunity. Nucleic Acids Res, 50, 6414–6422.

