## Supplementary figure for "ATP-dependent restriction enzyme SauUSI from *Staphylococcus aureus* is also a *bona fide* single strand DNA endonuclease"

### Slide 1
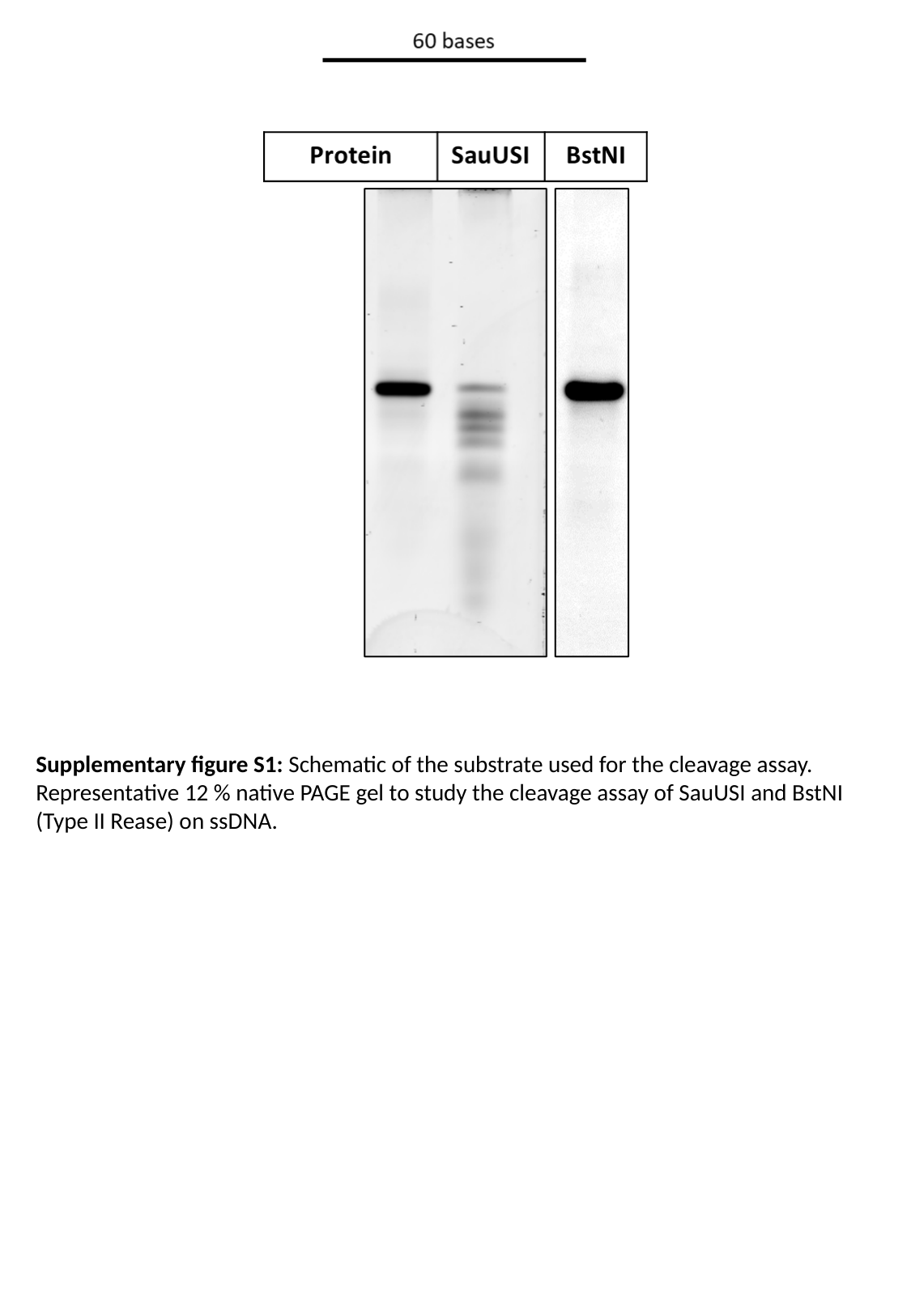

Supplementary figure S1: Schematic of the substrate used for the cleavage assay. Representative 12 % native PAGE gel to study the cleavage assay of SauUSI and BstNI (Type II Rease) on ssDNA.

### Slide 2
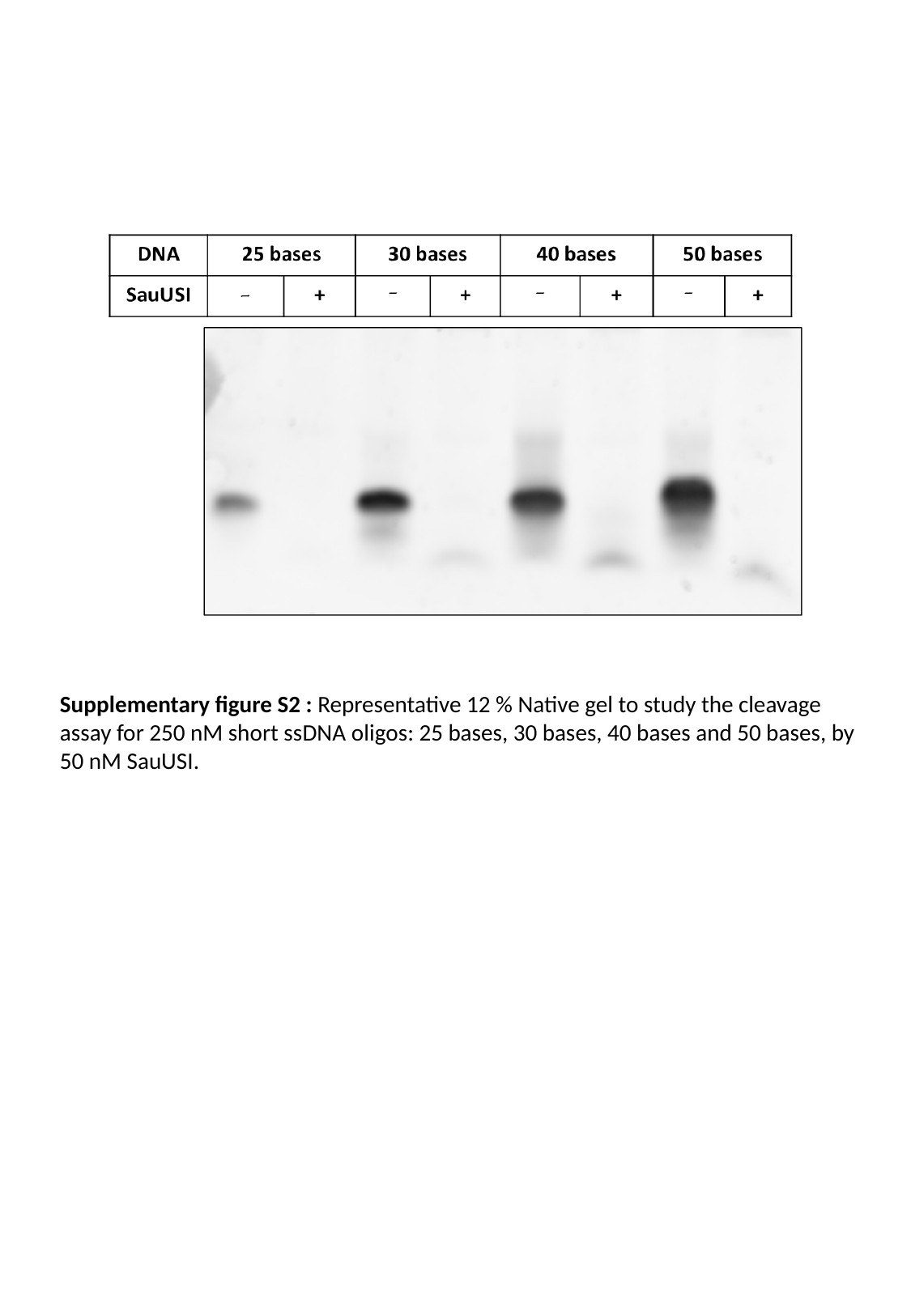

Supplementary figure S2 : Representative 12 % Native gel to study the cleavage assay for 250 nM short ssDNA oligos: 25 bases, 30 bases, 40 bases and 50 bases, by 50 nM SauUSI.

### Slide 3
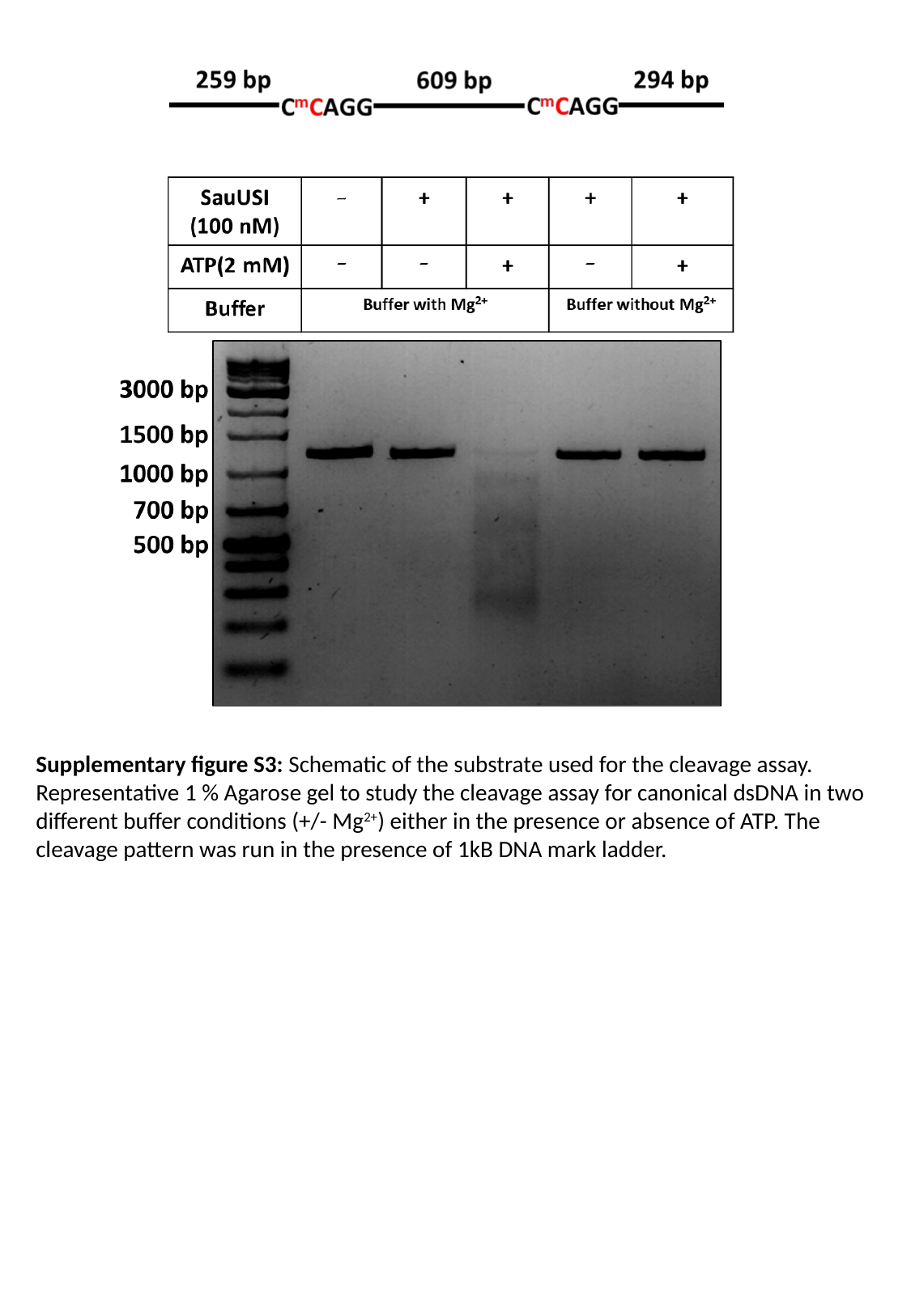

Supplementary figure S3: Schematic of the substrate used for the cleavage assay. Representative 1 % Agarose gel to study the cleavage assay for canonical dsDNA in two different buffer conditions (+/- Mg^2+^) either in the presence or absence of ATP. The cleavage pattern was run in the presence of 1kB DNA mark ladder.

### Slide 4
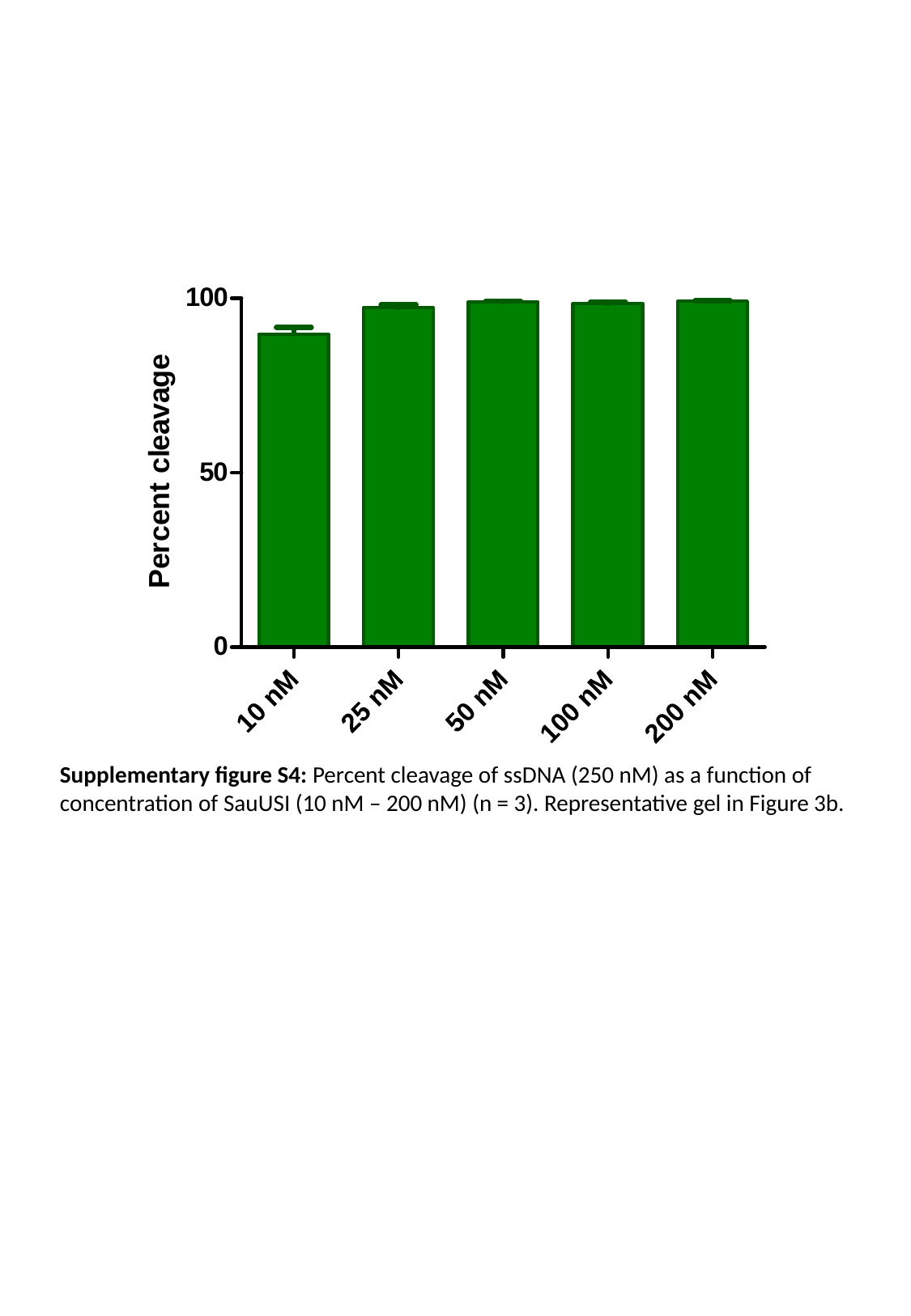

Supplementary figure S4: Percent cleavage of ssDNA (250 nM) as a function of concentration of SauUSI (10 nM – 200 nM) (n = 3). Representative gel in Figure 3b.

### Slide 5
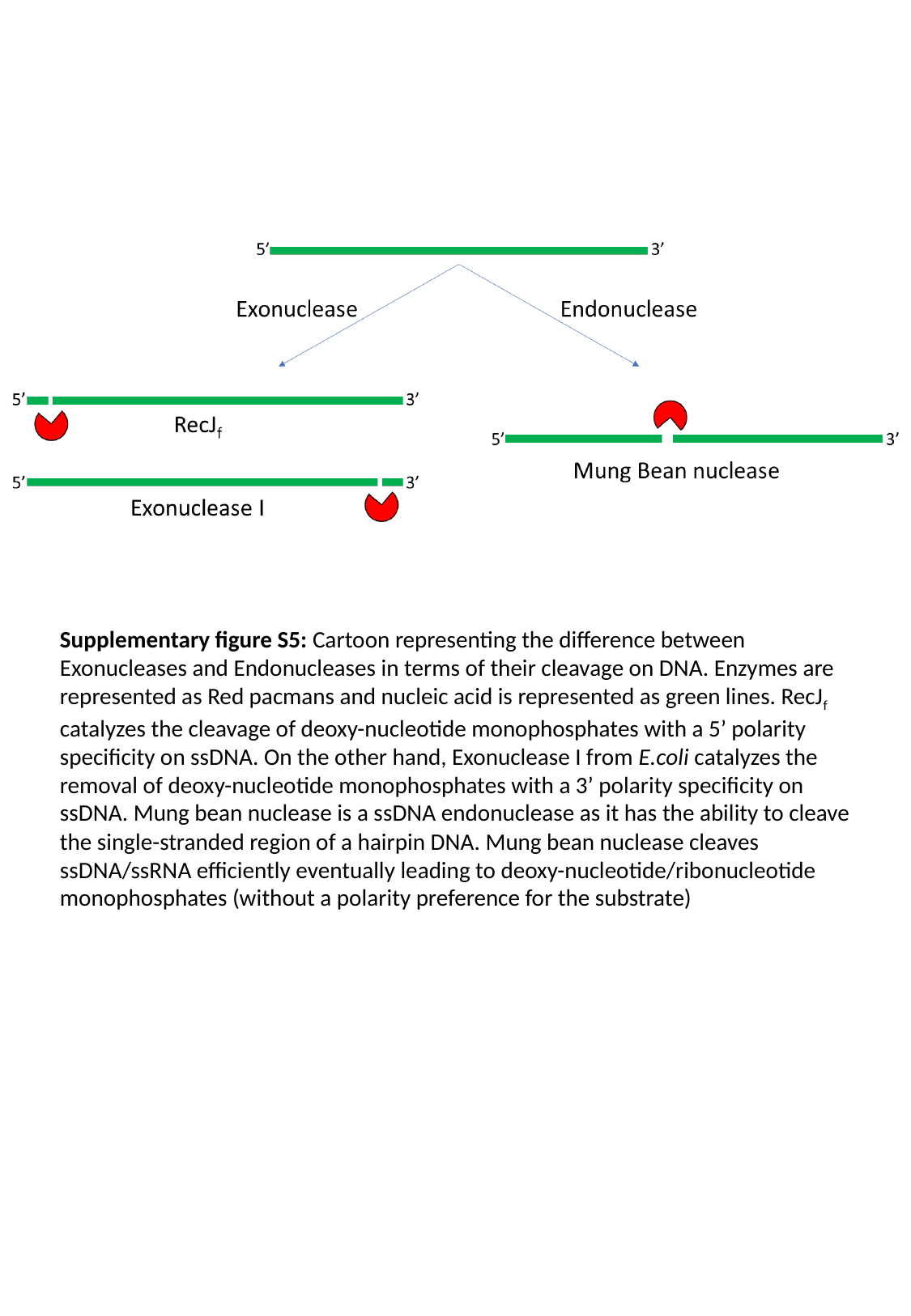

Supplementary figure S5: Cartoon representing the difference between Exonucleases and Endonucleases in terms of their cleavage on DNA. Enzymes are represented as Red pacmans and nucleic acid is represented as green lines. RecJ_f_ catalyzes the cleavage of deoxy-nucleotide monophosphates with a 5’ polarity specificity on ssDNA. On the other hand, Exonuclease I from E.coli catalyzes the removal of deoxy-nucleotide monophosphates with a 3’ polarity specificity on ssDNA. Mung bean nuclease is a ssDNA endonuclease as it has the ability to cleave the single-stranded region of a hairpin DNA. Mung bean nuclease cleaves ssDNA/ssRNA efficiently eventually leading to deoxy-nucleotide/ribonucleotide monophosphates (without a polarity preference for the substrate)

### Slide 6
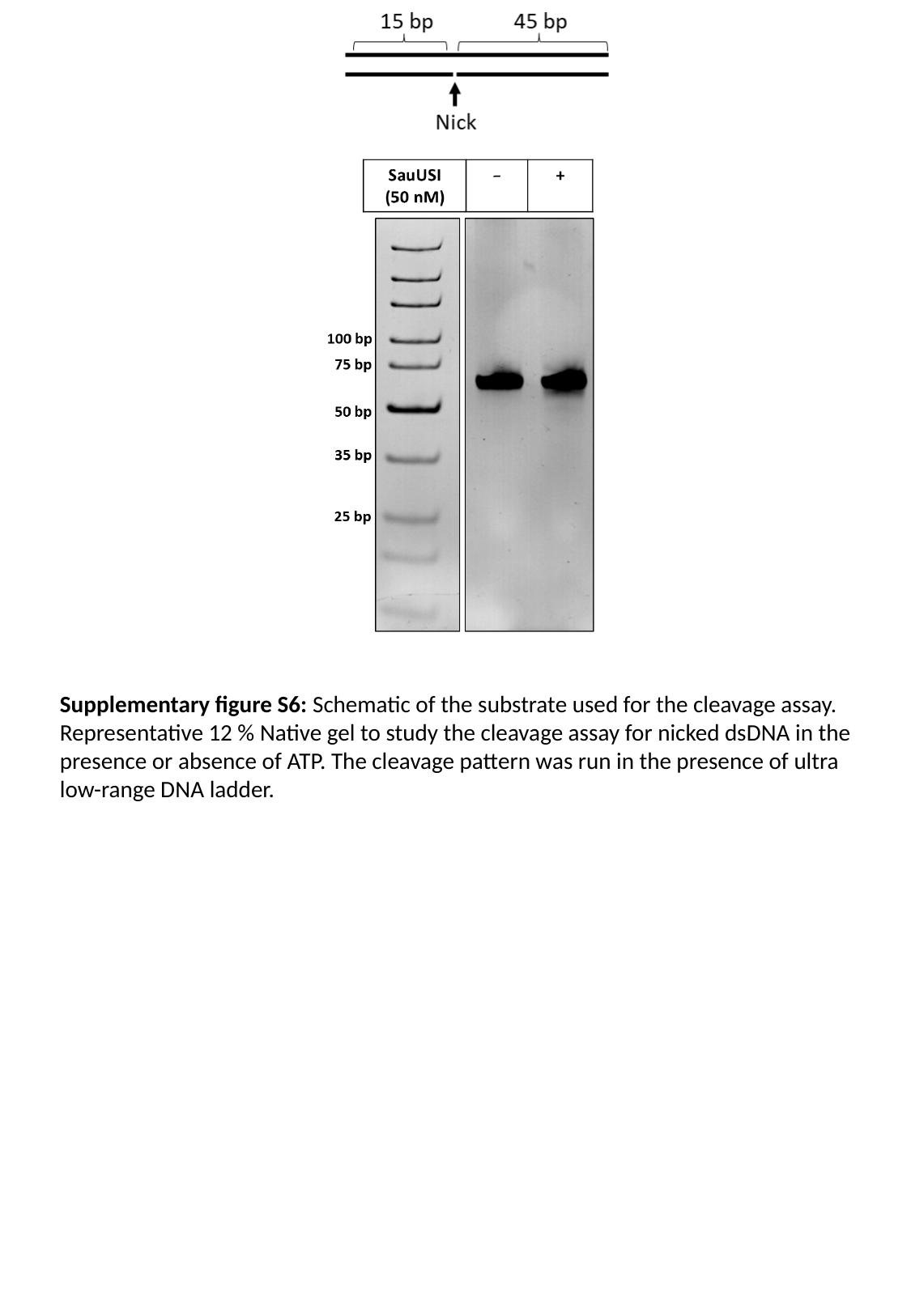

Supplementary figure S6: Schematic of the substrate used for the cleavage assay. Representative 12 % Native gel to study the cleavage assay for nicked dsDNA in the presence or absence of ATP. The cleavage pattern was run in the presence of ultra low-range DNA ladder.

### Slide 7
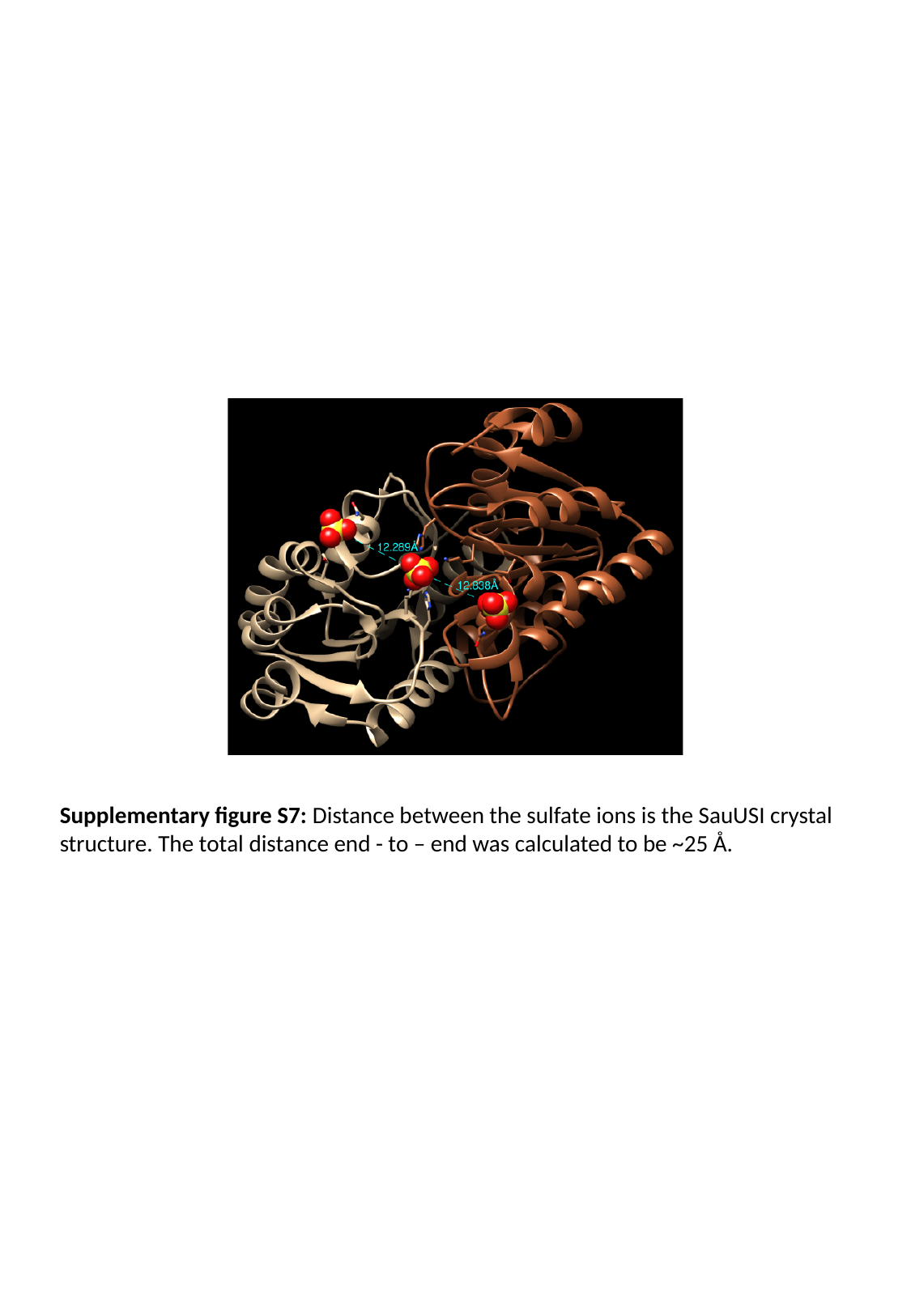

Supplementary figure S7: Distance between the sulfate ions is the SauUSI crystal structure. The total distance end - to – end was calculated to be ~25 Å.
