## Supplementary table for "ATP-dependent restriction enzyme SauUSI from *Staphylococcus aureus* is also a *bona fide* single strand DNA endonuclease"

| **Size (Bases) / Name** | **Substrate** | **Sequence (5' → 3')** |
|---|---|---|
| 60 | | • GCCGCGGATCCTTCACAAGACGATTACCGGCGCTGAAAGACCAGCATATGCAATTTGACT |
| 60 | | • AGTCAAATTGCATATGCTGGTCTTTCAGCGCCGGTAATCGTCTTGTGAAGGATCCGCGGC |
| 50 | | • ATGTCGTCACAAGCTCTTCACATCTATGCTAAATTCTCCTGGCGTAGTCC |
| 40 | | • GAAAGCGTTTGACGTGGACTACGTGGTCAATAGCACAACA |
| 30 | | • CATCTATGCTAAATTCTCCTGGCGTAGTCC |
| 25 | | • ATCCTTTGGCTTCGAATCCAGCAAT |
| 200 | | • ATGCTCGTCAGGGGGGCGGAGCCTATGGAAAAACGCCAGCAACGCGGCCTTTTTACGGTTC/5Med C/TGGCCTTTTGCTGGCCTTTTGCTCACATGTTCTTTCCTGCGTTATCCCCTGATCCTGGTCTGTGGATA ACCGTATTACCGCCTTTGAGTGAGCTGATACCGCTCGCCGCAGCCGAACGACCGAGCGCAGCGAGTC AGT |
| 36 | | • AGCAGAACGATGTGCCGCATCAAAAACAATATAATC |
| 20 (RNA) | | • GGGAUUAAUACGACUCACUG |
| Hairpin | 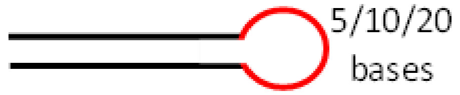  | • GCCCTTATTCCGATAGTGCTCTCTCTCTCTCTCTCTCACTATCGGAATAAGGGC<br>• GCCCTTATTCCGATAGTGCTCTCTCTCTCACTATCGGAATAAGGGC<br>• GCCCTTATTCCGATAGTGCTCTCCACTATCGGAATAAGGGC |
| Mismatch bubble | 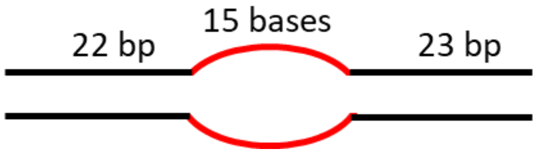  | • GCCGCGGATCCTTCACAAGACATTTTTTTTTTTTTTTAGACCAGCATATGCAATTTGACT<br>• AGTCAAATTGCATATGCTGGTCTCCCCCCCCCCCCCCCTGTCTTGTGAAGGATCCGCGGC |
| Substrate with 5' overhang | 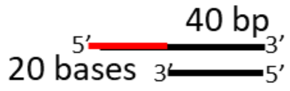  | • GCCGCGGATCCTTCACAAGACGATTACCGGCGCTGAAAGACCAGCATATGCAATTTGACT<br>• AGTCAAATTGCATATGCTGGTCTTTCAGCGCCGGTAATCG |
| Substrate with 3' overhang | 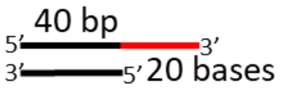  | • GCCGCGGATCCTTCACAAGACGATTACCGGCGCTGAAAGACCAGCATATGCAATTTGACT<br>• TCTTTCAGCGCCGGTAATCGTCTTGTGAAGGATCCGCGGC |
| Substrate with ssDNA gap | 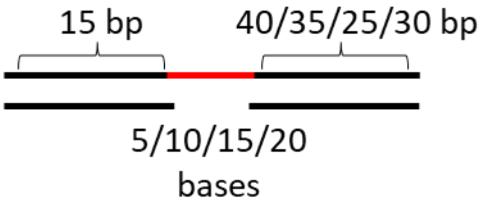  | • GCCGCGGATCCTTCACAAGACGATTACCGGCGCTGAAAGACCAGCATATGCAATTTGACT<br>• TGAAGGATCCGCGGC<br>• AGTCAAATTGCATATGCTGGTCTTT<br>• AGTCAAATTGCATATGCTGGTCTTTCAGCGCCGGT<br>• AGTCAAATTGCATATGCTGGTCTTTCAGCGCCGGTAATCG<br>• AGTCAAATTGCATATGCTGGTCTTTCAGCG |
